# Drivers of individual plant species contributions to *β*-diversity are scale-dependent

**DOI:** 10.1101/2025.08.11.669629

**Authors:** Rona Learmonth, Francis Q. Brearley, Adam Clark, Aaron S. David, Catalina Estrada, Fatih Fazlioglu, Ivan de Klee, Daniela Konrad, Lizzie Lemon, Olivia Millen, NJ Pates, Linda Pieber, Megan Sherlock, Elizabeth G. Simpson, Scott G. Ward, Hannah J. White, William D. Pearse

## Abstract

Species introductions and native extirpations are driving biotic homogenisation in plant communities by reducing *β*-diversity. Individual species vary in their contributions to *β*-diversity (Species Contribution to *β*-Diversity; species-*β*), yet our understanding of how species characteristics shape these contributions remains limited. Additionally, although the ecological processes influencing *β*-diversity are known to vary with spatial scale, we lack understanding of how species contributions, or their underlying determinants, change across scales. Here, we modelled how plant functional traits, phylogenetic relatedness, and introduction status influence their contributions to *β*-diversity using plant community data from 429 plots surveyed from 2017-2023 across three nested spatial scales (up to 1 km^2^) in nine sites spanning four countries. We extended the analysis to broader spatial extents (100 km^2^ to the entire UK) using GBIF occurrence data. We found that functional traits associated with competitive ability influenced species-*β*, but the direction and strength of their effects varied with scale. Likewise, phylogenetic novelty increased species-*β* at small scales but reduced it at larger ones. After accounting for traits and phylogeny, introduced species consistently contributed less to *β*-diversity than native species—especially at broader spatial extents. These results demonstrate that species’ ecological and evolutionary characteristics shape their contributions to *β*-diversity, but that these effects are scale-dependent. Our findings highlight the importance of scale-explicit approaches in understanding both the determinants of *β*-diversity, and how we can combat its loss to mitigate biotic homogenisation.

## 1 INTRODUCTION

CCurrently, biotic homogenisation, defined as increasing similarity or declining *β*-diversity of communities across space, is occurring at unprecedented rates (Petsch et al., 2022). Biotic homogenisation of plant communities has been reported across ecosystems (Bando et al., 2023; Britton et al., 2009; Daru et al., 2021; Pearse et al., 2018; Yang et al., 2015) and occurs due to local extinctions of native species and/or the increasing spread of non-native, often human-introduced species (Petsch et al., 2022). This increasing homogenisation is concerning, as declines in *β*-diversity are associated with reduced ecosystem services, stability (Fantinato et al., 2023), and multifunctionality (Hautier et al., 2018). Improving our knowledge of the underlying factors shaping patterns of *β*-diversity may allow us to better predict and mitigate biotic homogenisation (Olden et al., 2018).

Legendre & De Cáceres (2013) introduced a method for quantifying the role of individual species in shaping *β*-diversity: species contributions to *β*-diversity, hereafter referred to as species-*β*. This metric expresses how much a species contributes to the variation in species composition (*β*-diversity) between sites. Species that are highly widespread or dominant in all sites will typically contribute less, while those that vary greatly in presence or abundance between locations contribute more. This method allows us to move beyond whole-community measures and evaluate the homogenising or diversifying effects of specific species by quantifying the degree to which they each impact *β*-diversity. This method has been widely applied in plants and invertebrates, (da Silva et al., 2018; He et al., 2022; Heino & Grönroos, 2017; Pozzobom et al., 2020; Santos et al., 2021; Wang et al., 2023). However, the species characteristics, such as functional traits or phylogenetic relatedness which determine species-*β* remain uncertain. The Legendre & De Cáceres (2013) framework has been extended to identify individual species contributions to phylogenetic or functional *β*-diversity through matrices of phylogenetic and functional resemblance across species occurrence data (Nakamura et al., 2020). Our approach differs from this in that it assesses our capacity to predict individual species-*β* from functional traits and phylogenetic relatedness, instead of aiming to calculate functional or phylogenetic *β*-diversity themselves. Identifying traits linked to species-*β* could inform efforts to maintain *β*-diversity, for example, by prioritising the species expected to contribute most to *β*-diversity or controlling introduced species which contribute least.

Plants species’ functional traits influence their distribution via environmental trait filtering (Da et al., 2022), as well as their impacts on community structure, ecosystem properties, and interactions (Funk et al., 2017). Therefore, there have been calls for further research into the effects of plant traits on species-*β* (He et al., 2022). The potential role of ecological traits in influencing species-*β* is of particular interest in introduced species, whose rapid spread is among the greatest drivers of biotic homogenisation (Daru et al., 2021; Petsch et al., 2022). Non-native plant species with traits which confer competitive advantages tend to impact the diversity of native communities more negatively (Daly et al., 2023), so these traits may be associated with lower species-*β*. Greater relative height, lower specific leaf area (SLA), and lower seed mass are associated with increased competitiveness. These traits may enable competitive exclusion of other plant species (Feng et al., 2019; He et al., 2022) by promoting greater resource acquisition and invasiveness (Mathakutha et al., 2019; Wang et al., 2018). Additionally, some traits such as seed mass are known to differ between invasive and native populations, with greater seed mass in invasive populations thought to facilitate out-competition of native species (Daws et al., 2007). While the impact of functional traits on plants’ ability to drive biotic homogenisation has occasionally been tested (Stotz et al., 2019), the general relationships between plant functional traits and *β*-diversity remain uncertain. Addressing this knowledge gap is difficult because associations between functional traits and invasion success are context-dependent. For example, where taller introduced plants were shown to have a competitive advantage in grasslands (Stotz et al., 2019), harsh environmental filtering in a sub-Antarctic study resulted in outperformance by shorter introduced plants (Mathakutha et al., 2019). As a result, it remains unclear whether traits can reliably predict species-*β* for either native or introduced species across different environments.

As well as traits, a species’ phylogenetic distance from the rest of its community may influence its species-*β*. Using phylogenetic relatedness to infer species-*β* may offer an advantage over using functional trait values, as it is not limited by application of a mean trait value or assumptions of which traits are most ecologically relevant to competitive abilities in each context (Strauss et al., 2006). While the impact of phylogenetic relatedness on species-*β* remains untested, its role in mediating an introduced species’ effects on the wider community has frequently been investigated in the context of invasion success. However, studies disagree in whether they report greater invasion success for species that are more closely (Lim et al., 2014; Qian, 2023) or distantly (Feng et al., 2019) related to the native community.

It has been suggested that the inconsistent relationships between phylogenetic distance and species impacts on *β*-diversity in introduced species may be due to variation in the spatial scale assessed (Park et al., 2020). This is because studies at small spatial scales, at the ‘local community’ level, determined by dispersal limitations, detect competitive interactions between closely related introduced and native species (Vamosi et al., 2009). As the geographic distance increases, the outcome of competition is generally thought to become less important than environmental filtering in determining the success and impacts of invasive species (McGrannachan et al., 2020). In the context of species-*β*, this suggests that more closely related species may have lower species-*β* at small scales due to negative impacts of competition, but greater species-*β* at broader scales (Figure 1) due to the greater abundance and occupancy of phylogenetically clustered species which share similar environmental tolerances (He et al., 2022; Heino & Grönroos, 2017). Similarly, the influence of environmental filtering at large scales may obscure the impacts of functional traits which would be detected at small scales. Previously, the determinants of species-*β* have typically only been assessed at single scales, but to understand patterns of *β*-diversity across multiple scales, it is essential that we untangle how the complex drivers of species-*β* vary with scale (Figure 1). By investigating drivers across multiple nested spatial scales, we can improve our understanding of scale dependence in patterns of *β*-diversity (Crawley & Harral, 2001) and identify how conservation measures should be targeted at different spatial scales.

**FIGURE 1.**
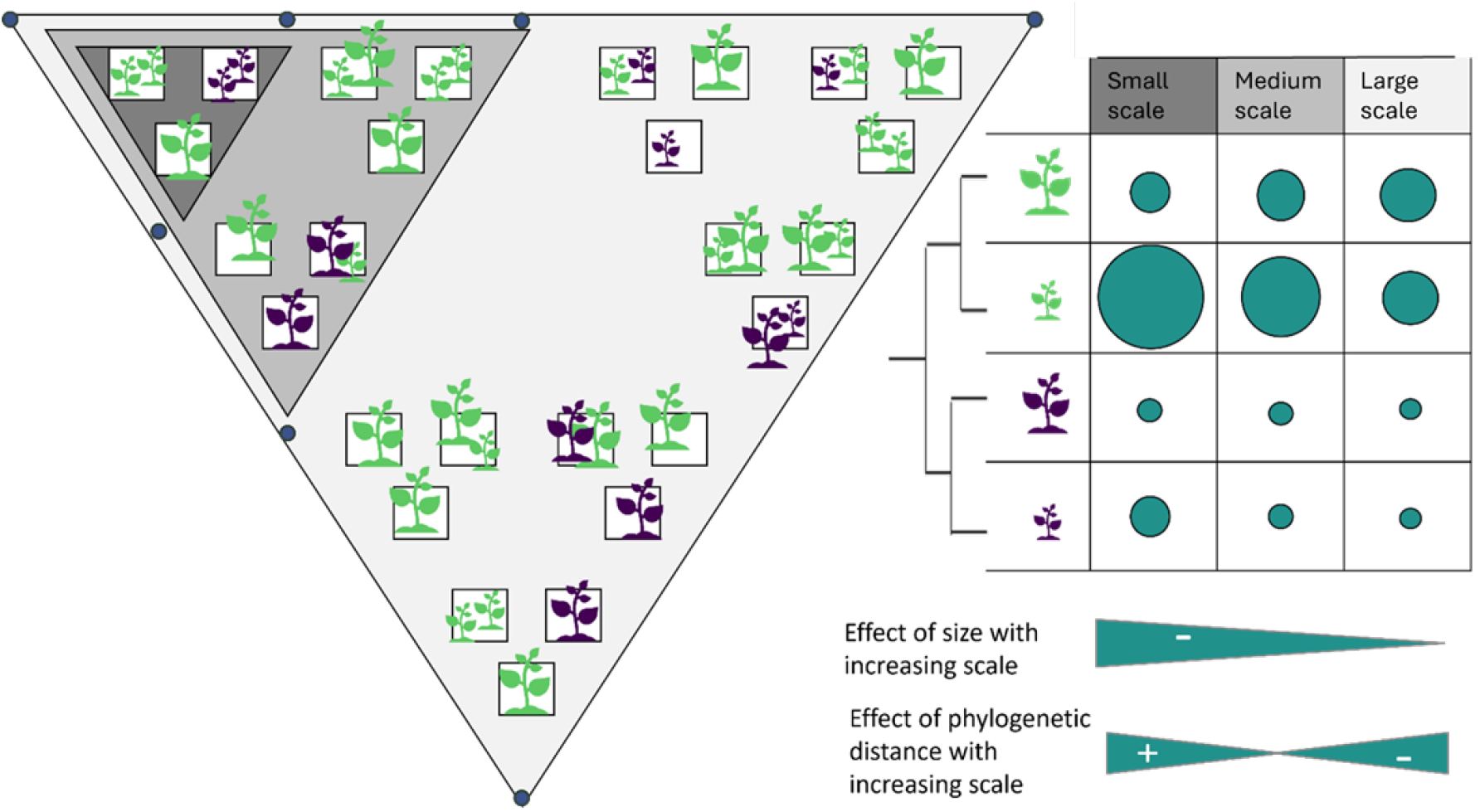
Conceptual diagram of how the determinants of species contributions to β-diversity may vary with spatial scale. On the left is a sampling schematic where white boxes indicate sampled plots, which are arranged in three levels of nested equilateral triangles to make up a site of 27 plots. Each of these sampling plots differs in its species composition: the plants within vary in terms of functional traits (size, as a proxy for competitiveness; shown by the size of the icons) and introduction status (colour; green for native and purple for introduced). The three grey triangles overlaid on the sampling schematic indicate the three levels at which species-β was calculated for each site. Species-β was calculated by comparing plots at the smallest spatial scale (minor triad; darkest grey), medium spatial scales (major triad; medium grey), and large scales (full fractal extent; lightest grey). This fractal sampling approach was also employed in Simpson & Pearse (2021). The blue circles indicate the arrangement of the minimal seven plots to be sampled at each fractal site in order to have at least three sites per spatial scale. Each species present is shown in the table on the right, with a phylogeny indicating the phylogenetic distance between them. This table shows the hypothesised effects of different species characteristics on their species-β. The species-β for each species at each of the three spatial scales is indicated by a blue dot, where larger dots indicate greater contributions, meaning the species explains more of the difference between the communities. species-β is influenced by species size at small scales, with larger species making smaller contributions at the patch and local levels, and this effect disappearing at the regional level. For example, the large native (green) species contributes less to β-diversity than the smaller native species at small scales, but their contributions are equal at large scales. This scale-dependent negative relationship between size and species-β is indicated by the arrow beneath the table. Species-β values are also influenced by the phylogenetic distance of the species, based on the branch lengths between them shown in the phylogeny. More distantly related species have greater species-β at small scales, but lower species-β at large scales, as shown by the arrow beneath the table. Lastly, introduced (purple) species make smaller contributions than native species across scales. For example, as in native species, the taller introduced species contributes less at small scales, with this effect disappearing at large scales. However, regardless of size, the introduced species make smaller relative contributions than the native species for each scale considered.

**FIGURE 2.**
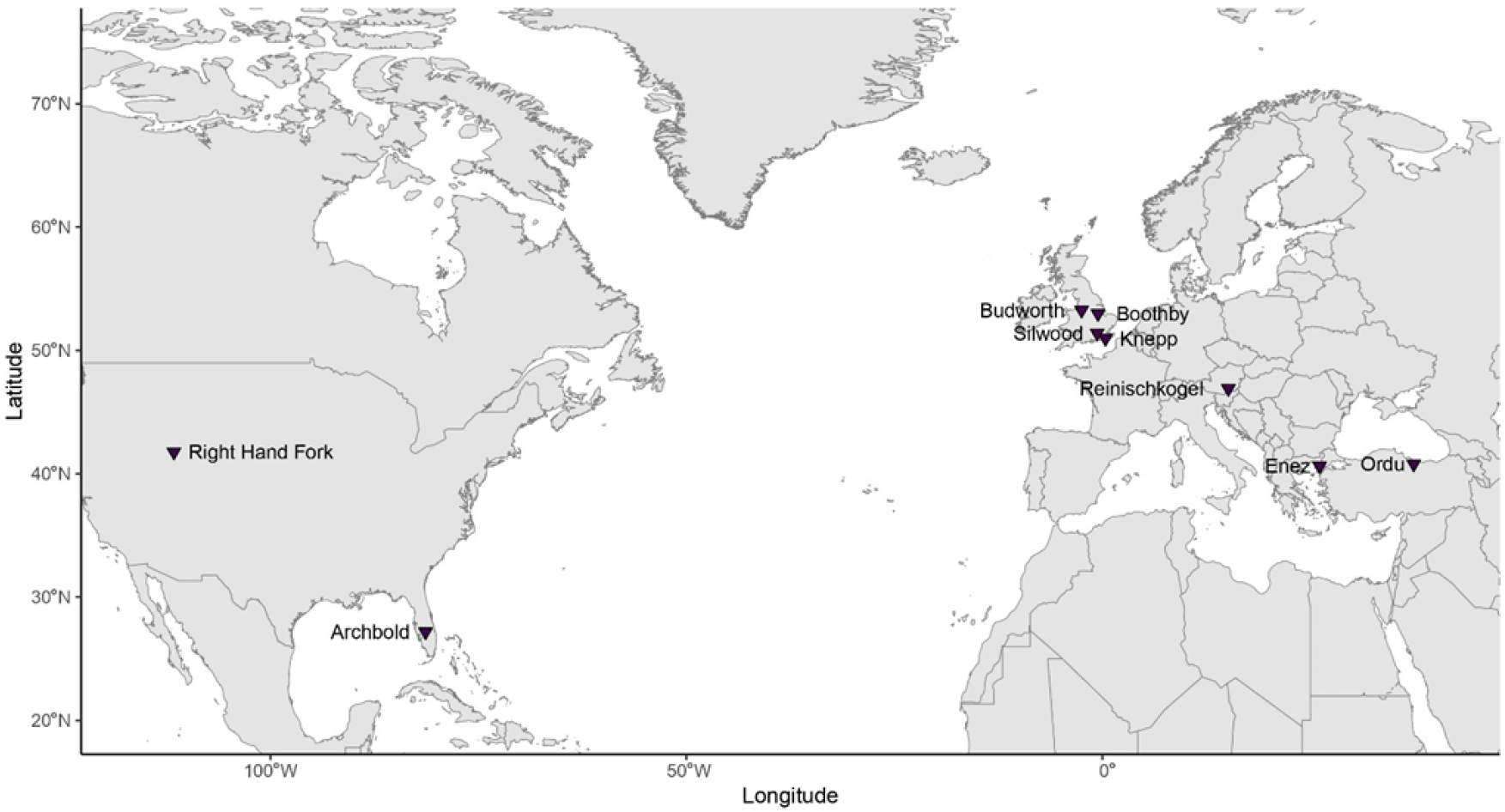
Fractal sampling site map. Triangles indicate the locations of the nine fractal sites surveyed in this study. Further information on the location, years sampled, and number of plots sampled, for each site can be found in Supplementary Materials Table 1.

Here, we use plant community data from the Ecological Fractal Network (EFN), which applies a nested fractal sampling design, to assess how functional traits, phylogenetic relatedness, and introduction status shape species-*β* across multiple spatial scales. This design allows us to test whether the ecological and evolutionary determinants of species-*β* shift from local to broad scales. We predicted that more competitive generalist species, which tend to be those with greater height, as well as lower SLA and seed mass (Funk et al., 2017; Daws et al., 2007), would contribute less to *β*-diversity, particularly at small scales. However, we predicted that environmental filtering would obscure the impacts of competitive interactions at larger spatial scales, resulting in declining importance of functional traits in assessing biotic homogenisation with increasing scale (McGrannachan et al., 2020). In addition, we expected species-*β* to increase at small scales but decline at large scales as phylogenetic distance increases since the impacts of competition at small scales become less influential than their shared environmental tolerances at large scales. To investigate the potential impacts of introduced species on biotic homogenisation, we also assessed whether native and introduced species varied in either their contributions to *β*-diversity or the relative importance of their determinants. We tested these predictions using plant community composition data collected at three spatial scales of nested fractal sampling across 429 plots in nine sites internationally. These smaller-scale data were supplemented with data extracted from the Global Biodiversity Information Facility (GBIF) for only the UK to consider two wider scales.

### 2 MATERIALS AND METHODS

We investigated the impacts of variation in three functional traits (SLA, seed mass, and height), the phylogenetic distance of plant species from others in their community, and introduction status on species contributions to *β*-diversity (species-*β*) across spatial scales. We used Ecological Fractal Network (EFN) data at three smaller scales, and GBIF data at larger scales from 10 km^2^ to the entire UK. EFN data was collected between 2017 and 2023 across nine sites, and is available, alongside all R scripts used, in the Supplementary Materials.

### 2.1 Plant community composition data collection

We collected plant community composition data from 429 unique 1 m^2^ plots across four countries between 2017 and 2023. The habitats surveyed across these sites included agricultural land, forests, and grasslands. We used a fractal sampling design made up of equilateral triangles nested at three spatial scales (see Figure 1). The overlapping vertices of the triangles at each spatial scale mean this design efficiently surveys the *β*-diversity of multiple spatial scales with equal coverage but minimal sampling effort (Marsh & Ewers, 2013). At each site, a minimum of seven plots, arranged as in Figure 1, of the full 27 making up each fractal were sampled (full site details are given in SI Table 1, Figure 1) to allow at least three sampled plots for each fractal scale. At all sites other than Right Hand Fork (see Supplementary Materials), the equilateral triangles had sides measuring 100 m for the small scale, 300 m for the medium scale and 900 m for the large scale (full fractal extent). At some sites (Knepp, Boothby, Right Hand Fork), sets of three fractal triangles were arranged into a larger equilateral triangle pattern. In these sites, each individual fractal was referred to as a ‘block’. We assessed the three scales individually for each block to make the scales assessed comparable across sites.

The plant community composition of each plot was measured by recording the percentage cover of all plant species present within each quadrant (0.5 × 0.5 m) of a 1 m^2^ quadrat. Quadrat placement was determined from mapped fractal layouts. Where sites were sampled in multiple years, quadrat placement was kept within 3 m of the previous year’s sampling. Where trees were present at the placement site, the quadrat was moved minimally such that it could be laid flat. Thus, only herbaceous and shrub-level plant cover was included, with overhanging trees not recorded. Only taxa identified to the species level were included in subsequent analyses. This sampling design is explained in full by Simpson and Pearse (2021) and is intended to maximise statistical power to detect shifts across environmental gradients and spatial scales. We recognise that our sampling is unevenly distributed with a bias to North America and Europe but, importantly, we account for this in our broader scale analysis accordingly (outlined below).

### 2.2 GBIF data collection

For the UK only, we assessed even larger scales, where we extracted abundance-weighted plant community composition data from GBIF using the R package *rgbif* (version 3.8.0, Chamberlain et al., 2023). We extracted occurrence data for a 100 km^2^ grid surrounding each of the fractal study sites (Figure 3a), hereafter referred to as the ‘Site’ level. For each site, we extracted the occurrence data of all observations in the Kingdom ‘Plantae’ with coordinate references for each year in which it was sampled in the fractal network. We determined the extent of each wider site based on its maximum and minimum latitude and longitude calculated from the coordinates of a central sampling point within the fractal site. We then divided the extracted occurrence data into 10 × 10 grids of 1 km^2^ grid cells, with the entirety of each fractal network site included in a single cell (Figure 3b). Similarly, for the largest level considered, we extracted all GBIF plant occurrence records in the United Kingdom for the year 2022, which was split into the 10 × 10 grid cells between which *β*-diversity was calculated and decomposed into species-*β* (Figure 3a). For these two broadest spatial scales, we chose to consider only GBIF data from the UK as sampling intensity was not constant around the world and this region was best covered by relatively consistent sampling effort surrounding the fractal sites. We conducted additional exploratory analyses to determine the impacts of considering international sites within the fractal network but only the UK at broader scales and found that analyses considering only UK data across all scales did not qualitatively differ from our findings including all sites. These data were accessed on 18/07/23.

**FIGURE 3.**
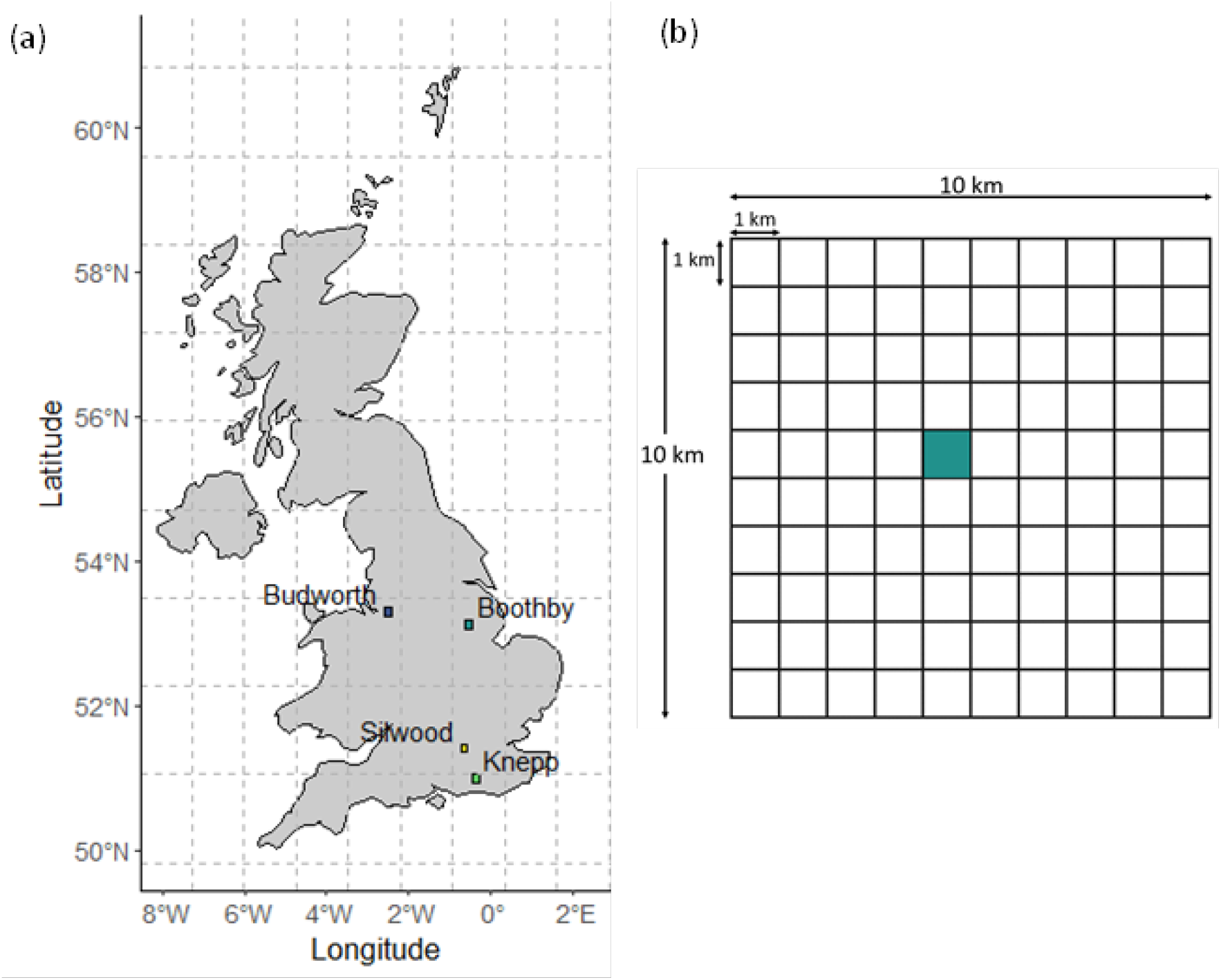
Sampling design for Site and UK level of analysis. (a) Map of the UK showing the locations of the four UK sites (Boothby Farm, Budworth, Knepp Estate and Silwood Park). Each site is indicated by a coloured box measuring 100 km^2^ centred on the 1 km^2^ area sampled within the fractal network. The overlaid grey dotted lines on the UK map indicate the 10 × 10 grid of boxes (A1-J10, approx. 135 km x 90 km) within which grid cells were considered as ‘plots’ for the whole UK level analyses. (b) Grid showing a zoomed in representation of a ‘Site’ comparable to the coloured boxes shown in Figure 3a. The full 10 × 10 grid of 1 km^2^ cells represents the whole site level, while the blue square indicates the central cell of the 1 km^2^ full fractal extent.

We conducted all data cleaning and analyses in R (version 4.3.1 R Core Team, 2023), with the same analyses applied to each spatial scale. While the sampling methodology of opportunistic GBIF data differs from the targeted sampling used at the fractal scales, expanding the scale of inquiry allows us to simultaneously assess local heterogeneity and broader turnover due to differing habitat types or climatic conditions. By applying identical analyses, we minimise the impact of the differing data sources to allow coarse comparison between spatial scales.

For each spatial scale, we classified species as ‘Native’ or ‘Introduced’ for each site in which they were found based on native/introduced range maps produced by the Royal Botanic Gardens, Kew (POWO, 2023). For some widespread species, this meant they were classified as ‘Native’ in some sites but ‘Introduced’ in others. While invasive species are accepted to be those which threaten native biodiversity, ecosystem functioning or have negative impacts on economies or public health, there are inconsistencies in how these criteria are addressed (Matthews et al., 2017). Therefore, introduced species were not subclassified into invasive and non-invasive species.

### 2.3 Trait data

We retrieved mean trait values for each species from the Botanical Information and Ecology Network (BIEN) using the function BIEN_trait_mean in the *BIEN* R package (version 1.2.6, Maitner, 2023). This function extracts trait values for each species, which are inferred from the genus or family mean when species-specific values are unavailable. Therefore, the conclusions drawn from this inferred data are conservative, as extreme trait values are averaged out at higher taxonomic levels. Using this method provided complete trait data for 503 unique species in the fractal sampling network, compared to 251 if species-specific values were used. Using this method also included all nine sampled sites, while using only species-specific values entirely excludes the Archbold, Florida site, which has a high proportion of endemic species. The traits used were the total plant height (m), log_10_ seed mass (mg) and specific leaf area (m^2^ kg^-1^), measured as leaf area (m^2^) per unit leaf dry mass (kg). We selected these traits as they are known correlates of stress tolerance, resource acquisition and competitiveness (Feng et al., 2019) which have been shown to relate to invasiveness (El-Barougy et al., 2021; Mathakutha et al., 2019) and were therefore thought likely to influence species-*β*.

### 2.4 Phylogenetic data

To investigate the impact of phylogenetic distance on species-*β*, we first pruned a large vascular plant phylogeny (Zanne et al., 2015) to produce phylogenies of the species present in each sampling site for each spatial scale studied, and filled in missing species using ‘congeneric.merge’ within the package *pez* (version 1.2-4, Pearse et al., 2015). We then converted these phylogenies to community data matrices using the package *picante* (version 1.8.2, Kembel et al., 2020). We used the function ‘cophenetic.phylo’ in the package *ape* (version 5.8, Paradis et al., 2023) to calculate the pairwise distances between all species in the community matrix based on the branch lengths of the phylogenetic tree. We then calculated mean phylogenetic distance (MPD) for each species (MSPD) from all others in its community, representing the mean evolutionary distance between them in terms of substitutions per sequence site.

We used a novel metric, community-standardised mean species pairwise distance (csMSPD) to calculate the relative phylogenetic distinctiveness of a species relative to its community. This differs from the standardised effect size of mean pairwise distance metric (SES_MPD_), which assesses MPD at the community level against a null model to test for non-random community structure (Kembel, 2009). Our approach is similar to the phylogenetic diversity field approach (Villalobos et al., 2013) but differs in that rather than phylogenetic species variability (PSV) and phylogenetic species clustering (PSC) indices, we use phylogenetic distance which gives readily interpretable units in terms of phylogenetic branch lengths.

This relative measure of each species’ phylogenetic distance from its community, csMSPD, was calculated for species i as its observed mean pairwise distance from its community (MSPD_i_) minus the mean pairwise distance (MPD) for the entire community, divided by the standard deviation of the community MPD.

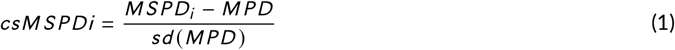

This standardised metric was calculated for each species at each scale, making phylogenetic distances comparable across all sites and scales. Here we emphasise that, while we are using phylogenetic metrics, we are using them to summarise species and not infer ecological assembly processes (see Mayfield & Levine, 2010). Where sites were sampled across multiple years, csMSPD was calculated separately for each year, as the community a given species may interact with varies in time as well as space.

### 2.5 Species contribution to *β*-diversity

To calculate *β*-diversity, we used Legendre & De Cáceres’ (2013) framework, as it allowed us to decompose total *β*-diversity (BD_total_) into individual species-*β* values. Following Legendre & Gallagher (2001), we first Hellinger-transformed the community matrices to generate Hellinger matrices (YHel) for each year of sampling at each site and spatial scale with columns as species and rows as plots. The BD_total_ of each matrix was calculated from its sum of squared deviations from its column means (SS_total_). Then we partitioned BD_total_ into the relative contribution of each species (species-*β*) by calculating the proportion of total variation (SS_total_) explained by species j (SS_j_):

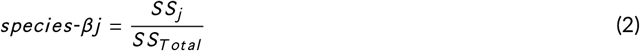

This calculation was performed using the function ‘beta.div’ in the package *adespatial* (version 0.3-23, Dray et al., 2023). The *β*-diversity was calculated and decomposed into individual species contributions at the small, medium and large scales in the fractal sites, and between grid cells within the ‘Site’ and ‘UK’ levels. Calculating values of species-*β* separately for each year of sampling in a given plot provided a context-dependent measure of species-*β* which could vary spatiotemporally. Although this method does result in pseudoreplication of species across sites, using mean values of csMSPD and species-*β* experienced by a species across multiple sites would obscure any impacts of species interactions in particular contexts.

### 2.6 Modelling species-*β* across spatial scales

As species-*β* varies between 0 and 1 and is beta-distributed, we used beta regression to model its determinants, as this accounted for the skew in the data and is the standard method of modelling species-*β* (da Silva et al., 2018; Santos et al., 2021; Tan et al., 2019; Xia et al., 2022). Using the R package *betareg* (version 3.1-4, Zeileis et al., 2021) with a logit link and maximum likelihood estimator, we modelled the determinants of species-*β* separately at each spatial scale. The explanatory factors considered were each species’ SLA, height, csMSPD, introduction status and the interactions of these species’ characteristics with introduction status. All trait values and phylogenetic distances were z-scaled prior to use in models to make regression coefficients comparable. In addition, we checked all model diagnostics and confirmed that they met the assumptions required.

We then conducted automated model selection and averaging in the package *MuMIn* (version 1.48.4, Bartoń, 2023) using the function ‘dredge’. Conditional average model coefficients were calculated as the mean of all models with AIC values within four units of the best model (delta () < 4) (Burnham & Anderson, 2004). This allowed us to untangle the determinants of species-*β* by testing whether functional traits or phylogenetic relatedness influenced species-*β*, and how these relationships differed between native and introduced species. Comparing the predictors of species-*β* across multiple spatial scales then allowed us to identify how spatial scale influences the relative importance of different drivers.

## 3 RESULTS

### 3.1 Summary statistics

We used data collected from nine sites (Figure 2, Supplementary Materials Table 1) from 2017-2023, comprising a total of 429 plots, 97 of which were sampled in at least two years, and included 585 species. GBIF data for the 100 km^2^ area surrounding each UK site included 221 grid cells and 428 species in total. At the UK level, GBIF observations were available for 63 grid cells, containing 3722 unique species. Of these, complete data were available for 503 species at fractal scales, and 1061 at GBIF scales, which were included in the final models. While the spatial scale of the Right Hand Fork site differed from all other sites, the results remain consistent when this site is excluded from models (see Supplementary Materials).

Across all data, mean plant height was 2.12 m (±0.03 SE) for natives and 1.96 m (±0.07 SE) for introduced species. Mean SLA was 25.2 m^2^ kg^-1^ (±0.06 SE) for natives and 25.9 m^2^ kg^-1^ (±0.11 SE) for introduced species. Mean seed mass was 103 mg (±4.2 SE) for natives and 318 mg (±25.6 SE) for introduced species. Lastly, the mean standardised effect size of the mean pairwise distance metric (csMSPD) was 0.023 (±0.005 SE) for natives and 0.125 (±0.017 SE) for introduced species.

### 3.2 Predicting species-*β* across spatial scales

The best models for predicting individual species contributions to *β*-diversity, here referred to as species-*β* at the small fractal scale 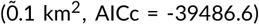, medium fractal scale 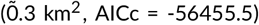, and large fractal scale 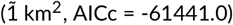 included all the variables. In each case, the interaction of seed mass and introduction status was the strongest predictor of species-*β*, but the relative importance of traits, phylogenetic distance, and introduction status varied (Figure 4). The best model for the site level (100 km^2^, AICc =-26423.3) likewise included all variables. However, at this scale, the mean model showed that the interaction of introduced status and SLA had greatest influence on species-*β* (estimate = 0.82 ±0.07 SE, z-value = 11.07, p<0.0001). At the UK level, the interaction of phylogenetic distance (csMSPD) with introduction status was excluded from the best model (AICc =-182711.5) but was included in the mean model (AICc = 1.5). At this scale, the strongest predictor of species-*β* was introduction status, with introduced species having lower species-*β* (estimate =-0.47 ±0.02 SE, z-value =-20.06, p<0.0001). The full model outputs are given in the Supplementary Information.

**FIGURE 4.**
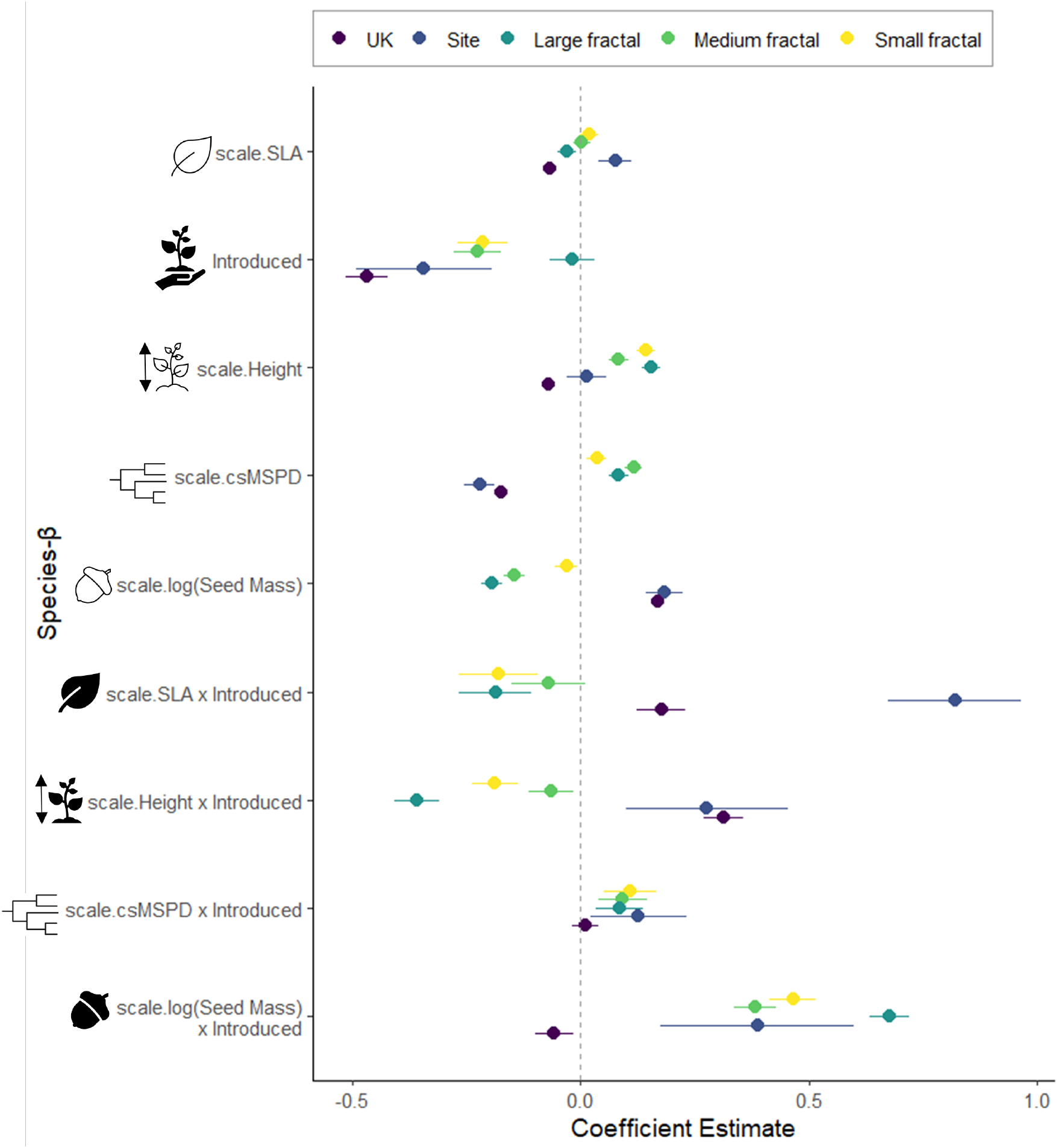
Effects of traits, phylogenetic distance, and introduction status across scales. Dot-whisker for each of the seven predictors shown on the vertical axis. The scaled height, csMSPD and SLA variables represent the effects of the driver in question on native species only, while the ‘x introduced’ drivers are interactions representing the contrast of the introduced to native value for that driver. The effect sizes from the mean models for the five spatial scales are indicated by different colours (purple to yellow; legend shown at the top of the figure). The coloured dots indicate the mean coefficient, with the lines showing their 95 % confidence intervals. Where these confidence intervals overlap with the 0 line, they are considered not to impact species-β. The sample size across scales varied between 11,633 individuals for the fractal levels (small, medium, large) to 16,580 individuals for the full UK extent. The direction of effects of the three functional traits (SLA, seed mass, and height) vary with spatial scale, with native and introduced species tending to show contrasting effects. Introduced species make lower contributions to β-diversity, particularly at large scales.

### 3.3 Effects of drivers of species-*β*

Against expectations that species with more competitive traits would have lower species-*β* and that this influence would be greatest at small scales, increased height was associated with higher species-*β* values in native plants at small scales, but lower at the UK-wide scale (Figure 4). For SLA, another trait associated with competitive ability (He et al., 2022), the direction of effect on native plants’ species-*β* was also scale dependent. However, in introduced species, greater SLA and height were linked to reduced species-*β* at the three smaller scales, as expected if these competitive traits allow them to competitively exclude other species from the local patch. Greater seed mass was associated with reduced species-*β* at the three smaller scales, and increased species-*β* at the two larger ones in native species. Again, introduced species showed a different trend, with greater seed mass associated with higher species-*β* at all scales but the UK-wide (Figure 4, 5). Contrary to expectations that the influence of traits would decrease with increasing spatial scale due to the greater effects of environmental filtering, there was no consistent change in the magnitude of the impact of any trait with increasing spatial scale, though the direction of the effect frequently differed between the fractal and GBIF scales.

**FIGURE 5.**
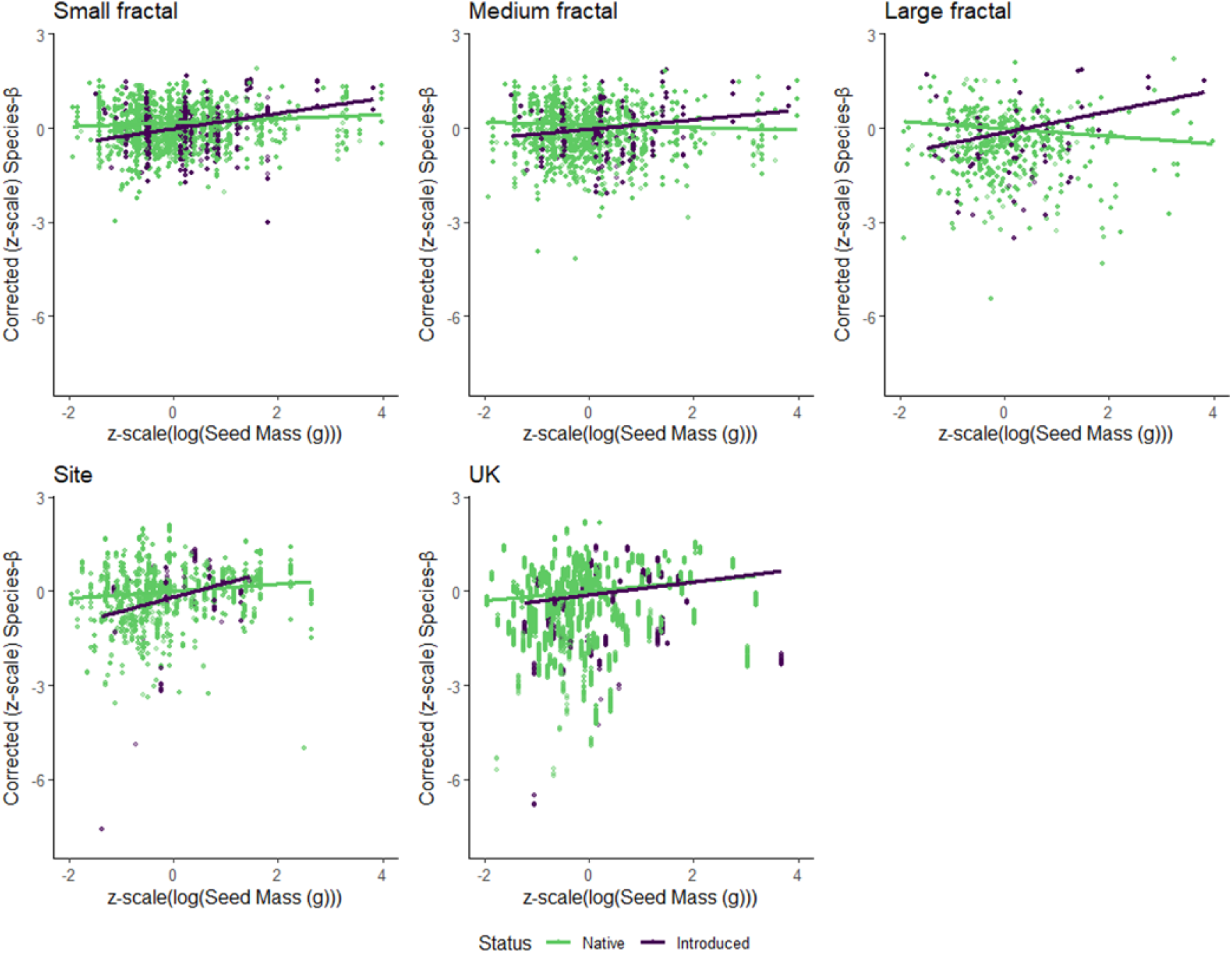
Effect of plant seed mass on species-β across five spatial scales. Partial beta regression plots showing corrected species’ contribution to *β*-diversity (species-*β*) plotted on the vertical axis against-scaled log seed mass (mg) for each of the five spatial scales modelled, plotted on the horizontal axis in size order from smallest (small fractal) to largest (‘UK’). The corrected values of species-*β* in each case represent the residuals of a beta regression model for species-*β* at the given scale which included all factors considered in Figure 4 except seed mass and its interaction with introduction status. These residuals represent the value of species-*β* when controlled for the other drivers considered. On each plot, the line indicates the fitted linear model. Native species are shown in green, while introduced species are shown in purple. In both native and introduced species, the direction of effect of seed mass on contribution to *β*-diversity depended on the scale of assessment.

Native plants’ species-*β* increased with increasing phylogenetic distance (csMSPD) of the focal native species from its community at the small fractal scales, but decreased at larger scales. Meanwhile in introduced species, species-*β* tended to increase with phylogenetic distance at all scales except the entire UK where there was no significant relationship.

## 4 DISCUSSION

Decomposing measures of *β*-diversity into the individual contributions of each species present, referred to here as species-*β*, could allow us to improve our understanding of the factors shaping patterns of diversity across scales. Our study works towards these goals by assessing the determinants of plant species contributions to *β*-diversity (species-*β*) across five nested spatial scales in terms of the impacts of plants’ functional traits, relatedness, and introduction status. We identified that plants’ species-*β* s are influenced by both their traits and relatedness, with the direction of these relationships differing between native and introduced species, and across scales.

### 4.1 Effects of functional traits on native species-*β* are scale-dependent

We found that all three functional traits considered (SLA, seed mass, and height) were significant predictors of species-*β* across multiple spatial scales. While previous studies have consistently reported abundance and occupancy as drivers of species-*β* (He et al., 2022; Heino & Grönroos, 2017; da Silva et al., 2018; Santos et al., 2021; Xia et al., 2022), the few investigations of the impact of functional traits have produced contrasting results. Some previous studies have reported relationships between functional distinctiveness, a composite measure of functional traits, and plant species-*β* (Pozzobom et al., 2020; Wang et al., 2023), while others have reported no influence of traits (Tan et al., 2019). We found that trait effects on species-*β* change with scale, which may explain inconsistent findings in previous work conducted at different scales. For example, SLA may increase, decrease, or have no effect on native plants’ species-*β* depending on the spatial scale addressed. This highlights the need for more multi-scale studies to identify how the scale of enquiry influences findings and identify reliable indicators across scales. Being able to predict species-*β* from indicators such as widely available functional trait values would allow us to more effectively target conservation efforts towards species likely to contribute most to *β*-diversity (Tan et al., 2019; Wang et al., 2023). However, we show here that simple trait-based indicators would not be reliable unless scale is accounted for.

### 4.2 Functional traits impact species-*β* differently in introduced and native species

Height affected species-*β* of introduced plants differently depending on scale. At the three smaller fractal scales, introduced species-*β* decreased with increasing plant height, but at wider scales showed slight increases with plant height. The declines in species-*β* with increasing height are consistent with the idea that taller plants are stronger competitors which outcompete shorter ones (El-Barougy et al., 2021; Hejda et al., 2009; Wang et al., 2018) and so contribute less to *β*-diversity. Additionally, as in our study, these effects of competition are generally detected more strongly at small scales. Thus, our findings suggest competitive taller introduced species can reduce *β*-diversity and thereby drive biotic homogenisation (Stotz et al., 2019). The increases in species-*β* with plant height at broader scales are consistent with the slight increases in species-*β* with increasing diameter at breast height (DBH), another proxy for plant size, previously reported in a broad-scale study in deciduous forests in China (He et al., 2022). These associations may occur because broad-scale studies capture multiple habitat types, thereby masking the context-dependent impacts of traits (Daly et al., 2023).

Seed mass is determined by a trade-off between competitive and colonisation capacity. Heavier seeds tend to be better able to survive and compete, while lighter seeds tend to disperse further and allow colonisation (Turnbull et al., 2004), suggesting we might expect competitive advantages of larger seeds at small scales, and smaller seeds at large scales. The finding that species-*β* increased at small scales but reduced at large scales with increasing seed mass in native species is therefore surprising. In introduced species, on the other hand, species-*β* increased with seed mass across all scales. Together, these findings may reflect shifts in the trade-offs between dispersal and competitiveness that depend on context, scale and introduction status. We also found inconsistent effects of SLA on both native and introduced species-*β*, against predictions of increased competitiveness with larger SLA (Feng et al., 2019). Together, these findings show that functional traits may significantly impact species-*β*, but that their scale-dependent effects mean they are likely not reliable indicators of individual species impacts on *β*-diversity.

However, the limitations of our findings on the effect of traits on species-*β* should be noted. Firstly, while these traits did not have a consistent effect across spatial scales, it is possible that other biological traits such as dispersal mode (Tan et al., 2019), ecological traits such as niche position (da Silva et al., 2018), or composite trait metrics such as functional distinctiveness (Wang et al., 2023) may better correlate with species-*β* across scales. Secondly, the trait values we used were extracted from the BIEN trait database (Maitner, 2023), meaning they represent the mean value of at least one botanical observation. However, plant functional traits show high levels of intraspecific variation (Siefert et al., 2015) so the species trait values used – sometimes based on a single observation – are a limited reflection of the actual traits of the plants surveyed. This does not fully represent the difference, for example, between maximum height and the achieved height of sampled plants, resulting in inflated estimates of height. In addition, the ratio of intra-to interspecific trait variation declines with increasing spatial scale (Da et al., 2022; Siefert et al., 2015). This suggests that intraspecific variation may more strongly obscure effects of traits on species-*β* at smaller scales, so the mean effect sizes here may indicate a conservative estimate of actual trait effects at small scales. Future studies should thus use trait measurements taken directly from the plants sampled to account for this influence of intraspecific trait variation.

### 4.3 Closer relatives contribute more to species-*β* at small scales, but less at large scales

As well as functional traits, we found that a species’ phylogenetic distance from its community influenced species-*β*. In native plants, species which were more distantly related to their community had slightly greater species-*β*. This matches the expectation that closely related species would compete at small scales, potentially driving competitive exclusion and reduced contributions to *β*-diversity (Cavender-Bares et al., 2006). At wider scales, species-*β* declined with increasing phylogenetic distance, meaning focal species more closely related to their communities had greater species-*β* than more distant relatives. This may reflect a shift from competitive outcomes most strongly determining species abundance and occupancy across sites at small scales towards environmental filtering based on shared tolerances of closer relatives at larger scales (Park et al., 2020; Qian, 2023). With a greater increase at small scales, and a decrease at large scales, introduced species showed broadly similar patterns in species-*β* with increasing phylogenetic distance from the community. As expected if this pattern was driven by differences in competition intensity, species-*β* showed greater increases with phylogenetic distance at smaller spatial scales (Park et al., 2020; Vamosi et al., 2009). Pouteau et al. (2023) found a similar relationship between phylogenetic relatedness and impacts of introduced species, reporting that native plants’ extinction risk was greater when exposed to more closely related introduced plant species. Our finding indicates that phylogenetic distance plays a role in determining introduced species’ contributions to *β*-diversity, and therefore their impacts on patterns of diversity. This provides further support for the use of phylogenetic relatedness in predicting diversity impacts and prioritising the control of introduced species more closely related to the native community (Strauss et al., 2006).

### 4.4 Introduced species contribute less to *β*-diversity across spatial scales

Introduced species contributed less than native species at all scales except the large fractal scale, where there was no significant difference. The difference between native and introduced plant species-*β* tended to increase with spatial scale. This is partially consistent with the idea of biotic differentiation, where at small scales, the presence of introduced species can sometimes increase *β*-diversity as they increase the patchiness of the area (Danneyrolles et al., 2021; Martin & Wilsey, 2015). Both biotic homogenisation and biotic differentiation are scale-dependent processes, with differentiation thought to occur at small sampling scales, and homogenisation detected at wider grains (Qian & Qian, 2022). Evidence of this pattern has been shown empirically, with a meta-analysis demonstrating that detection of biotic homogenisation of tropical forests increased with the spatial scale of the study (Kramer et al., 2023). The majority of studies of the impacts of introduced plant species are conducted at small scales (Olden et al., 2018) which suggests they may underestimate the consequences of introduced species invasion for *β*-diversity (Bando et al., 2023). This strong scale-dependence of biotic homogenisation (Soares et al., 2019; Yang et al., 2015) highlights the need to address knowledge gaps regarding the scales at which introduced species impact *β*-diversity (Martin & Wilsey, 2015). This will allow us to generate stronger predictions of biotic homogenisation and prioritise the prevention and control of introduced species expected to contribute least to *β*-diversity.

Further research into the role of biotic invasion in species-level diversity impacts is now needed. A key next step would be to assess the impacts of introduction status beyond the binary ‘introduced’ and ‘native’ categories used here to assess differences between invasive and non-invasive species. This is important as invasive and non-invasive plants have previously been shown to differ in their mean trait values (Mathakutha et al., 2019), and phylogenetic distances from native communities (Strauss et al., 2006). Additionally, it has been shown that the role of phylogenetic relatedness in determining the success or consequences of an invasion varies with the stage of invasion studied (Omer et al., 2022; Qian, 2023), so it follows that their contributions to *β*-diversity may also vary with the stage of invasion. For example, biotic differentiation often occurs at early phases of invasion, which later gives way to biotic homogenisation as introduced species become widespread (Qian & Qian, 2022). As we were unable to account for invasion stage, the differences between native and widespread invasive plants’ contributions to *β*-diversity may exceed those reported here. Future studies addressing how the determinants of species-*β* vary with invasion could help us to better identify threats to *β*-diversity and devise frameworks for early detection of potentially harmful introduced species.

### 4.5 Conclusion

In conclusion, both functional traits and the phylogenetic relatedness of plant species impact their individual contributions to *β*-diversity, based on data across multiple spatial scales and countries. The variation in the relative importance of these drivers across spatial scales highlights the need for more spatially explicit studies of species-*β*. Working towards a better understanding of how individual species contribute to patterns of *β*-diversity, and the spatial scales on which these mechanisms act, is a key step in developing frameworks to understand how conservation efforts can best be targeted to preserve *β*-diversity against the current background of rapid global biotic homogenisation. Understanding how native and introduced species differ in their individual contributions to *β*-diversity offers a novel way to predict which introductions may pose greater threats to *β*-diversity and inform our management priorities.

## Supporting information

All supplemental information

## Supporting Information

All data used for this article are provided in support of this manuscript at: doi.org/10.5281/zenodo.15044233

## 6 ACKNOWLEDGEMENTS

WDP and the Pearse lab are supported by UKRI BB/Y008766/1, UKRI NE/X00547X/1, UKRI NE/X013022/1, the Alan Turing Institute FA.04, and the Singapore Green Finance Centre.

## Notes

https://doi.org/10.5281/zenodo.16793184

### Competing Interest Statement

Ivan de Klee is Head of Natural Capital, and Lizzie Lemon is Site & Communities Coordinator at Nattergal, which manages Boothby Wildland.
The remaining authors certify that they have no affiliations with, or involvement in, any organisation or entity with any financial or non-financial interest in the subject matter or materials discussed in this manuscript. We confirm that this work is original and is not under consideration for publication elsewhere.

