## Supplementary material for "Drivers of individual plant species contributions to *β*-diversity are scale-dependent": All supplemental information

**Supplementary Materials**

***Supplementary Materials Table 1:* Site details.** *Names, locations, habitat descriptions, and number of plots included from each year of sampling for the nine fractal network sites included in this study. Blocks indicate the number of individual three-level fractal triangles sampled at a site. Total spatial extent sampled indicates the length of the full fractal extent, and the full length across blocks (longest side of triangle).*

| **Site name** | **Country** | **Site coordinates** | **Brief description** | **Plots sampled per site** | **Total spatial extent sampled** |
| --- | --- | --- | --- | --- | --- |
| Archbold, Florida | USA | 27.1827° N,  -81.3520° W | Shrubland comprised of oak scrub, flatwoods, and seasonal ponds. | 2021: 27 plots | 900 m; N/A |
| Boothby | UK | 52.9943° N,  -0.5514° W | Agricultural land converted to rewilding in 2022. Crop fields, field-margins, occasional woodlands, and meadows. | 2022: 30 plots (3 blocks)  2023: 62 plots (3 blocks) | 900 m; 2700 m |
| Budworth | UK | 53.2719° N,  -2.5165° W | Deciduous woodland and heathland. | 2021: 7 plots | 900 m; N/A |
| Enez | Türkiye | 40.6395° N,  26.0778° E | Deciduous forest. | 2021: 7 plots | 900 m; N/A |
| Knepp Estate | UK | 50.9834° N,  -0.3547° W | Former agricultural land converted to rewilding between 2001-2006. Open woodland and grassland. | 2022: 32 plots (5 blocks)  2023: 73 plots (5 blocks) | 900 m; 1800 m |
| Ordu | Türkiye | 40.9714° N,  37.9645° E | Urban area mainly covered with hazelnut orchards. | 2021: 7 plots | 900 m; N/A |
| Reinischkogel | Austria | 46.9163° N,  15.1161° E | Mountainous region dominated by dense deciduous woodland with occasional grassland clearings. | 2021: 27 plots | 900 m; N/A |
| Right Hand Fork, Utah | USA | 41.7699° N,  -111.5850° W | Variable vegetation across a gradient defined by slope- aspect (Simpson & Pearse, 2021). | 2017: 27 plots (1 block)  2018: 78 plots (3 blocks)  2019: 27 plots (3 blocks)  2020: 27 plots (3 blocks)  2021: 27 plots (3 blocks) | 663 m; 1900 m |
| Silwood Park | UK | 51.4092° N,  -0.6415° W | Grassland and deciduous woodland, with occasional cultivated areas. | 2021: 2 plots  2022: 24 plots  2023: 21 plots | 900 m; N/A |

***Supplementary Materials Figure 1: Fractal sampling design schematics.*** *(a) The three levels of sampling within the fractal network, with the number of sites required for each level reported below. Grey triangles indicate the equilateral triangle addressed, while the green dots indicate the actual sampling plots. Adapted from Simpson and Pearse (2021). (b) Design and layout of sampling plots at each site, arranged as three levels of nested equilateral triangles within a 1 km^2^ area. The required plots, which were sampled as a minimum across all nine sites in each year in which they were sampled (1-4 years), are shown as green circles, with the optional plots shown as grey circles. (c) The quadrat (1 m^2^) representing each plot, indicated by the green circle surrounding the quadrat, which was split into four quadrants (0.25 m^2^), numbered as shown, oriented facing upslope.*

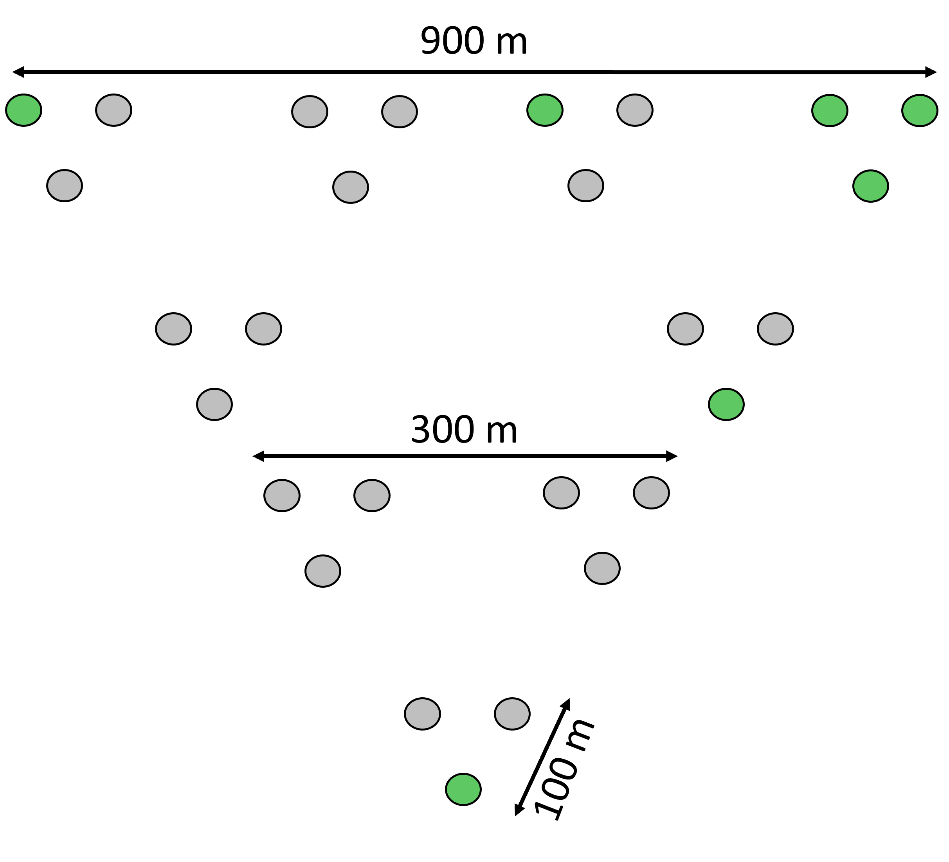

(a)

(b)

(c)

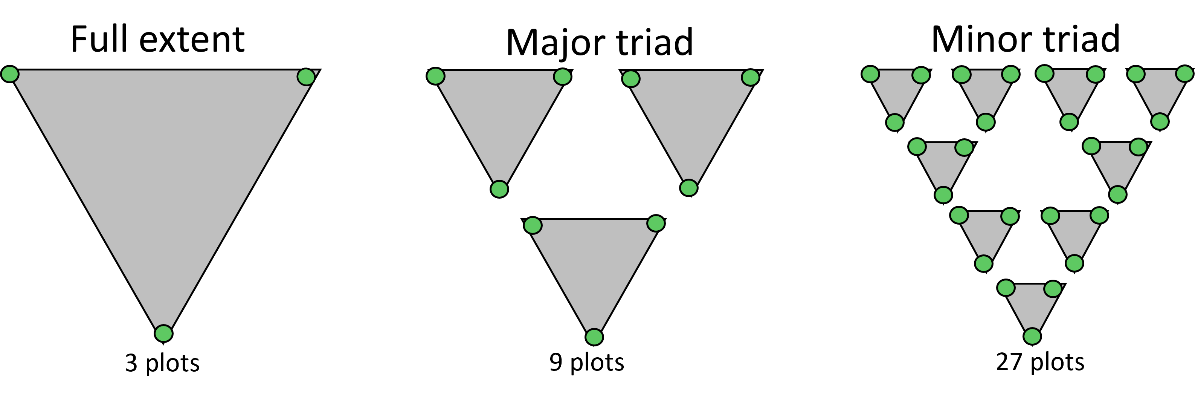

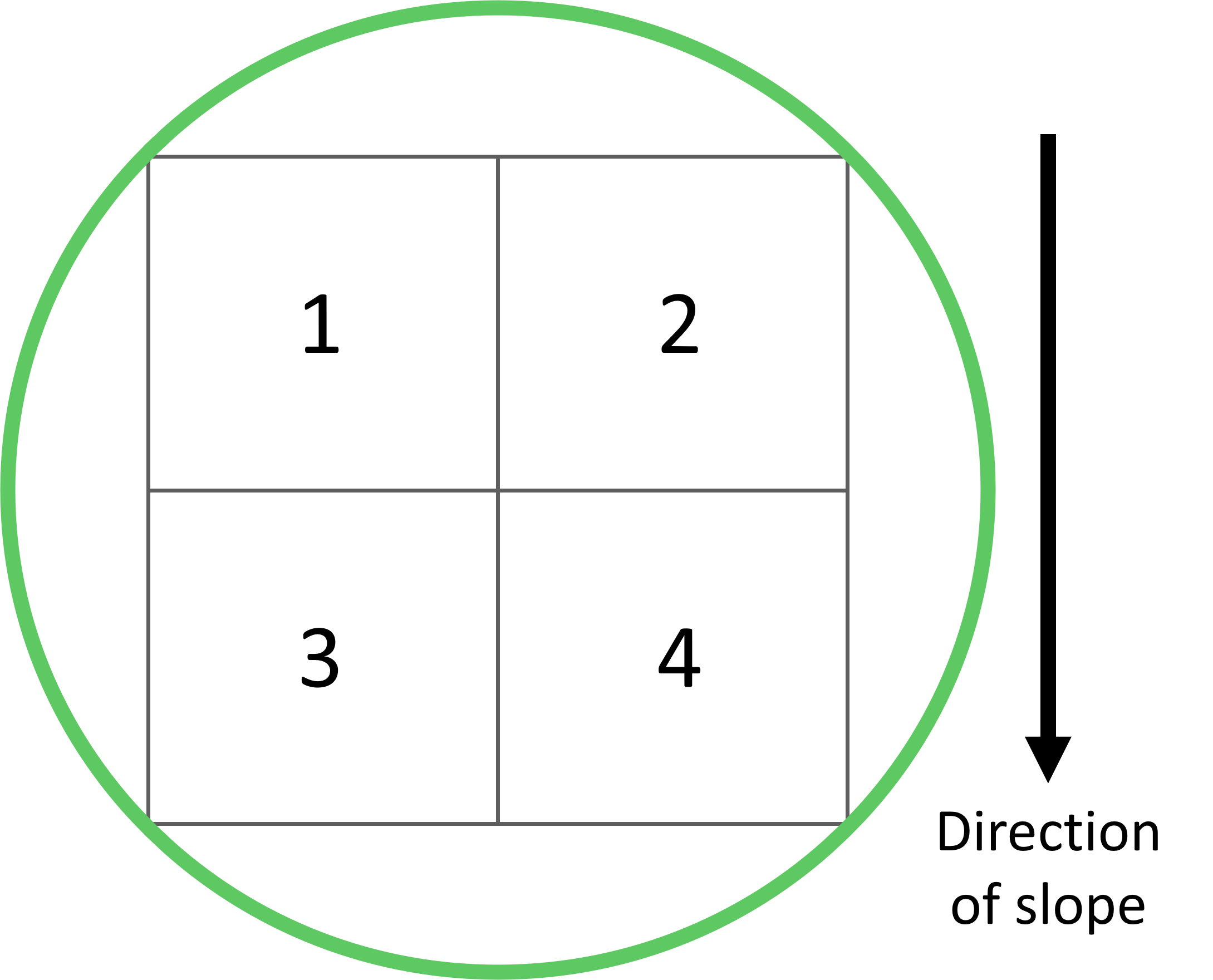

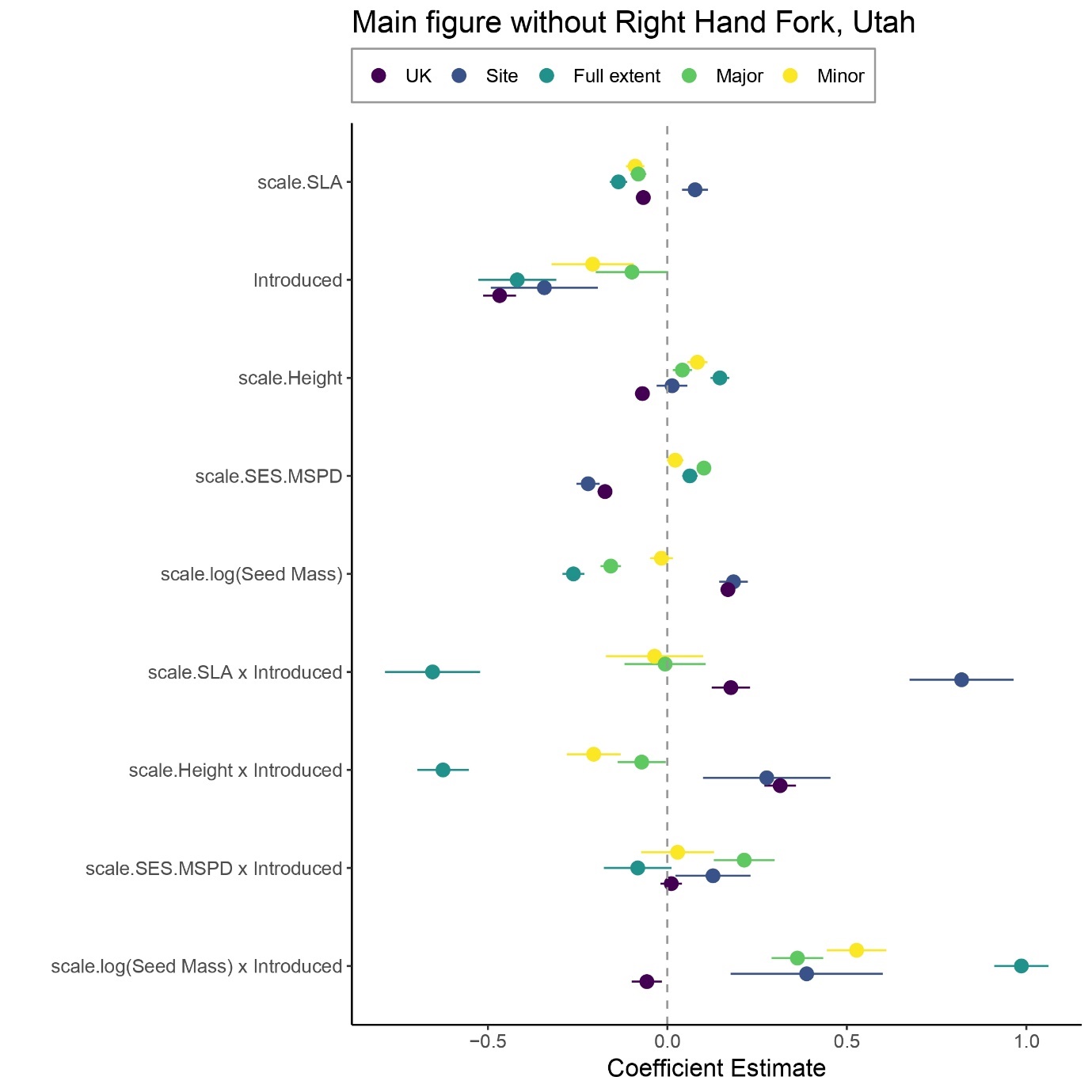

***Supplementary Materials Figure 2: Effects of traits, phylogenetic distance, and introduction status across scales excluding the site Right Hand Fork (RHF), Utah, USA.*** *The RHF fractal site differed slightly from all others in its dimensions, see Simpson & Pearse (2021) for full description. To test whether this difference in scale influenced the findings at fractal scales, the full analysis was repeated excluding this site. The above dot-whisker plot shows the estimated standardised effect size of the mean model on the horizontal axis plotted for each of the nine predictors shown on the vertical axis and differs from Figure 4 only in that its models excluded Right Hand Fork. This demonstrates that the difference in this fractal site’s size had limited qualitative impact.*

**Beta regression MuMIn selected model outputs**

***Supplementary Materials Table 2: Plot level model averaged coefficients (full average).*** *At the plot level, the best model contained all variables and was not averaged as all alternative models had ΔAICc > 4.*

| **Variable** | **Estimate** | **Standard error** | **z-value** | **p-value** |
| --- | --- | --- | --- | --- |
| SLA | 0.020 | 0.011 | 1.82 | 0.069 |
| Introduced | -0.214 | 0.028 | -7.73 | > 0.0001 |
| Height | 0.144 | 0.011 | 13.28 | > 0.0001 |
| SES_MSPD_ | 0.036 | 0.011 | 3.14 | 0.002 |
| Log(Seed mass) | -0.030 | 0.012 | -2.44 | 0.015 |
| SLA x Introduced | -0.178 | 0.044 | -4.05 | > 0.0001 |
| Height x Introduced | -0.187 | 0.026 | -7.15 | > 0.0001 |
| SES_MSPD_ x Introduced | 0.108 | 0.029 | 3.68 | 0.0002 |
| Log(Seed mass) x Introduced | 0.466 | 0.026 | 18.17 | > 0.0001 |

***Supplementary Materials Table 2: Minor triad model averaged coefficients (full average).*** *Model averages were made up of four component models, where one contained all explanatory variables except SLA and SLA x Introduced (df = 9, logLik = 28236.8, AICc = -56455.6, delta 0.00, weight = 0.46), one with all variables (df = 11, logLik = 28238.1, AICc = -56454.1, delta 1.50, weight = 0.22), one with all variables except SLA x Introduced (df = 10, logLik = 28236.8, AICc = -56453.6, delta 2.00, weight = 0.17), and lastly one not containing SLA, SLA x Introduced, and Height x Introduced (df = 8, logLik = 28234.7, AICc = -56453.4, delta 2.16, weight = 0.16).*

| **Variable** | **Estimate** | **Standard error** | **z-value** | **p-value** |
| --- | --- | --- | --- | --- |
| SLA | 0.004 | 0.010 | 0.37 | 0.715 |
| Introduced | -0.226 | 0.026 | -8.67 | < 0.0001 |
| Height | 0.084 | 0.011 | 7.89 | < 0.0001 |
| SES_MSPD_ | 0.117 | 0.010 | 12.05 | < 0.0001 |
| Log(Seed mass) | -0.145 | 0.011 | -12.85 | < 0.0001 |
| SLA x Introduced | -0.069 | 0.041 | -1.70 | 0.089 |
| Height x Introduced | -0.065 | 0.025 | -2.54 | 0.011 |
| SES_MSPD_ x Introduced | 0.093 | 0.027 | 3.40 | 0.001 |
| Log(Seed mass) x Introduced | 0.384 | 0.024 | 16.19 | < 0.0001 |

***Supplementary Materials Table 3: Major triad model averaged coefficients (full average).*** *At the major triad level, the best model contained all variables and was not averaged as all alternative models had ΔAICc > 4.*

| **Variable** | **Estimate** | **Standard error** | **z-value** | **p-value** |
| --- | --- | --- | --- | --- |
| SLA | -0.030 | 0.010 | -2.95 | 0.003 |
| Introduced | -0.017 | 0.025 | -0.69 | 0.489 |
| Height | 0.155 | 0.010 | 15.20 | < 0.0001 |
| SES_MSPD_ | 0.084 | 0.011 | 8.00 | < 0.0001 |
| Log(Seed mass) | -0.194 | 0.012 | -16.61 | < 0.0001 |
| SLA x Introduced | -0.187 | 0.040 | -4.61 | < 0.0001 |
| Height x Introduced | -0.358 | 0.025 | -14.52 | < 0.0001 |
| SES_MSPD_ x Introduced | 0.085 | 0.026 | 3.22 | 0.001 |
| Log(Seed mass) x Introduced | 0.677 | 0.022 | 30.51 | < 0.0001 |

***Supplementary Materials Table 4: Wider site area (100 km^2^) model averaged coefficients (full average).*** *Model averages were made up of two component models within delta 4 of each other, where one contained all variables (df = 11, logLik = 13222.7,, AICc = -26423.3, delta 0.00, weight = 0.72), and the other excluded only SES_MSPD_ x Introduced (df = 10, logLik = 13220.8,, AICc = -26421.5, delta 1.84, weight = 0.28),*

| **Variable** | **Estimate** | **Standard error** | **z-value** | **p-value** |
| --- | --- | --- | --- | --- |
| SLA | 0.077 | 0.018 | 4.19 | > 0.0001 |
| Introduced | -0.343 | 0.076 | -4.51 | > 0.0001 |
| Height | 0.013 | 0.022 | 0.60 | 0.549 |
| SES_MSPD_ | -0.221 | 0.017 | -13.33 | > 0.0001 |
| Log(Seed mass) | 0.184 | 0.020 | 9.05 | > 0.0001 |
| SLA x Introduced | 0.820 | 0.074 | 11.07 | > 0.0001 |
| Height x Introduced | 0.277 | 0.091 | 3.05 | 0.002 |
| SES_MSPD_ x Introduced | 0.127 | 0.053 | 2.38 | 0.0003 |
| Log(Seed mass) x Introduced | 0.388 | 0.108 | 3.59 | 0.017 |

***Supplementary Materials Table 5: Whole UK model averaged coefficients (full average).*** *Model averages were made up of two component models, where one contained all explanatory variables except SES_MSPD_ x Introduced (df = 10, logLik = 91365.8, AICc = -182711.5, delta = 0.00, weight = 0.68), and the other included all variables (df = 11, logLik = 91366.0, AICc = -182710.0, delta = 1.50, weight = 0.32).*

| **Variable** | **Estimate** | **Standard error** | **z-value** | **p-value** |
| --- | --- | --- | --- | --- |
| SLA | -0.067 | 0.006 | -11.05 | > 0.0001 |
| Introduced | -0.467 | 0.023 | -20.05 | > 0.0001 |
| Height | -0.069 | 0.007 | -9.99 | > 0.0001 |
| SES_MSPD_ | -0.174 | 0.007 | -26.21 | > 0.0001 |
| Log(Seed mass) | 0.169 | 0.007 | 25.18 | > 0.0001 |
| SLA x Introduced | 0.177 | 0.027 | 6.51 | > 0.0001 |
| Height x Introduced | 0.314 | 0.022 | 14.04 | > 0.0001 |
| SES_MSPD_ x Introduced | 0.011 | 0.015 | 0.71 | 0.476 |
| Log(Seed mass) x Introduced | -0.057 | 0.021 | -2.67 | 0.008 |

**BIEN trait data references**

Aakala, T., Shimatani, I., Abe, T., Kubota, Y. & Kuuluvainen, T. (2015) Data from: Crown asymmetry in high latitude forests: disentangling the directional effects of tree competition and solar radiation. *Oikos*. doi:[doi:10.5061/dryad.6t6gp](https://doi.org/doi:10.5061/dryad.6t6gp).

Abakumova, M., Zobel, K., Lepik, A. & Semchenko, M. (2016) Data from: Plasticity in plant functional traits is shaped by variability in neighbourhood species composition. *New Phytologist*. doi:[doi:10.5061/dryad.83g9k](https://doi.org/doi:10.5061/dryad.83g9k).

Ackerly, D.D. (2004) Adaptation, niche conservatism, and convergence: comparative studies of leaf evolution in the California chaparral. *The American Naturalist*. 163 (5), 654–671.

Ameztegui, A., Paquette, A., Shipley, B., Heym, M., Messier, C. & Gravel, D. (2016) Data from: Shade tolerance and the functional trait - demography relationship in temperate and boreal forests. *Functional Ecology*. doi:[doi:10.5061/dryad.12b0h](https://doi.org/doi:10.5061/dryad.12b0h).

Anderson-Teixeira, K., McGarvey, J., Muller-Landau, H., Park, J., Gonzalez-Akre, E., Herrmann, V., Bennett, A., So, C., Bourg, N., Thompson, J., McMahon, S. & McShea, W. (2015) Data from: Size-related scaling of tree form and function in a mixed-age forest. *Functional Ecology*. doi:[doi:10.5061/dryad.6nc8c](https://doi.org/doi:10.5061/dryad.6nc8c).

Anon (2013) *Forest Inventory and Analysis National Program*. <http://www.fia.fs.fed.us/>.

Bhaskar, R., Dawson, T. & Balvanera, P. (2014) Data from: Community assembly and functional diversity along succession post-management. *Functional Ecology*. doi:[doi:10.5061/dryad.6p9v5](https://doi.org/doi:10.5061/dryad.6p9v5).

Blonder, B., Buzzard, V., Simova, I., Sloat, L., Boyle, B., Lipson, R., Aguilar-Beaucage, B., Andrade, A., Barber, B., Barnes, C., & others (2012) The leaf-area shrinkage effect can bias paleoclimate and ecology research. *American Journal of Botany*. 99 (11), 1756–1763.

Boyero, L., Pearson, R., Hui, C., Gessner, M., Perez, J., et al. (2016) Data from: Biotic and abiotic variables influencing plant litter breakdown in streams: a global study. *Proceedings of the Royal Society B*. doi:[doi:10.5061/dryad.jg8r0](https://doi.org/doi:10.5061/dryad.jg8r0).

Bufford, J., Lurie, M. & Daehler, C. (2015) Data from: Biotic resistance to tropical ornamental invasion. *Journal of Ecology*. doi:[doi:10.5061/dryad.b1v2c](https://doi.org/doi:10.5061/dryad.b1v2c).

Burns, J., Halpern, S. & Winn, A. (2006) Data from: A test for a cost of opportunism in invasive species in the Commelinaceae. *Biological Invasions*. doi:[doi:10.5061/dryad.8107q](https://doi.org/doi:10.5061/dryad.8107q).

Carmona, C., Rota, C., Azcárate, F. & Peco, B. (2014) Data from: More for less: sampling strategies of plant functional traits across local environmental gradients. *Functional Ecology*. doi:[doi:10.5061/dryad.53550](https://doi.org/doi:10.5061/dryad.53550).

Carus, J., Paul, M. & Schröder, B. (2016) Data from: Vegetation as self-adaptive coastal protection: reduction of current velocity and morphologic plasticity of a brackish marsh pioneer. *Ecology and Evolution*. doi:[doi:10.5061/dryad.np6b8](https://doi.org/doi:10.5061/dryad.np6b8).

Cavender-Bares, J., González-Rodríguez, A., Eaton, D., Hipp, A., Beulke, A. & Manos, P. (2015) Data from: Phylogeny and biogeography of the American live oaks (Quercus subsection Virentes): a genomic and population genetics approach. *Molecular Ecology*. doi:[doi:10.5061/dryad.855pg](https://doi.org/doi:10.5061/dryad.855pg).

Cornwell, W.K., Schwilk, D.W. & Ackerly, D.D. (2006) A trait-based test for habitat filtering: convex hull volume. *Ecology*. 87 (6), 1465–1471.

Correia, M., Montesinos, D., French, K. & RodrA-guez-EcheverrA-a, S. (2016) Data from: Evidence for enemy release and increased seed production and size for two invasive Australian acacias. *Journal of Ecology*. doi:[doi:10.5061/dryad.f1kc3](https://doi.org/doi:10.5061/dryad.f1kc3).

Dalponte, M. & Coomes, D. (2016) Data from: Tree-centric mapping of forest carbon density from airborne laser scanning and hyperspectral data. *Methods in Ecology and Evolution*. doi:[doi:10.5061/dryad.hf5rh](https://doi.org/doi:10.5061/dryad.hf5rh).

de, E., la, Riva, PACrez-Ramos, I., Tosto, A., Navarro-FernA!ndez, C., Olmo, M., MaraA?A3n, T. & Villar, R. (2015) Data from: Disentangling the relative importance of species occurrence, abundance and intraspecific variability in community assembly: a trait-based approach at the whole-plant level in Mediterranean forests. *Oikos*. doi:[doi:10.5061/dryad.dr275.2](https://doi.org/doi:10.5061/dryad.dr275.2).

DeWalt, S.J., Bourdy, G., Chavez de Michel, L.R. & Quenevo, C. (1999) Ethnobotany of the Tacana: Quantitative inventories of two permanent plots of Northwestern Bolivia. *Economic Botany*. 53 (3), 237–260. doi:[10.1007/BF02866635](https://doi.org/10.1007/BF02866635).

Dostál, P., Fischer, M., Chytrý, M. & Prati, D. (2016) Data from: No evidence for larger leaf trait plasticity in ecological generalists compared to specialists. *Journal of Biogeography*. doi:[doi:10.5061/dryad.p3057](https://doi.org/doi:10.5061/dryad.p3057).

Easdale, T.A. & Healey, J.R. (2009) Resource-use-related traits correlate with population turnover rates, but not stem diameter growth rates, in 29 subtropical montane tree species. *Perspectives in Plant Ecology, Evolution and Systematics*. 11 (3), 203–218. doi:<http://dx.doi.org/10.1016/j.ppees.2009.03.001>.

Edwards, E., Chatelet, D., Sack, L. & Donoghue, M. (2014) Data from: Leaf lifespan and the leaf economic spectrum in the context of whole plant architecture. *Journal of Ecology*. doi:[doi:10.5061/dryad.61g42](https://doi.org/doi:10.5061/dryad.61g42).

Enquist, B.J., Sandel, B., Boyle, B., Svenning, J.-C., McGill, B.J., et al. (n.d.) *Botanical big data shows that plant diversity in the New World is driven by climatic-linked differences in evolutionary rates and biotic exclusion*.

Evans, L., Kaluthota, S., Pearce, D., Allan, G., Floate, K., Rood, S. & Whitham, T. (2016) Data from: Bud phenology and growth are subject to divergent selection across a latitudinal gradient in Populus angustifolia and impact adaptation across the distributional range and associated arthropods. *Ecology and Evolution*. doi:[doi:10.5061/dryad.ch720](https://doi.org/doi:10.5061/dryad.ch720).

Feng, Y. & van Kleunen, M. (2016) Data from: Phylogenetic and functional mechanisms of direct and indirect interactions among alien and native plants. *Journal of Ecology*. doi:[doi:10.5061/dryad.0672g](https://doi.org/doi:10.5061/dryad.0672g).

Fricke, E. & Wright, S. (2016) Data from: The mechanical defence advantage of small seeds. *Ecology Letters*. doi:[doi:10.5061/dryad.90f03](https://doi.org/doi:10.5061/dryad.90f03).

Gapare, W. (2015) Data from: Genetic parameters in subtropical pine F1 hybrids: heritabilities, between-trait correlations and genotype-by-environment interactions. *Tree Genetics & Genomes*. doi:[doi:10.5061/dryad.d5672](https://doi.org/doi:10.5061/dryad.d5672).

Gavinet, J., Prévosto, B. & Fernandez, C. (2016) Data from: Introducing resprouters to enhance Mediterranean forest resilience: importance of functional traits to select species according to a gradient of pine density. *Journal of Applied Ecology*. doi:[doi:10.5061/dryad.k67j5](https://doi.org/doi:10.5061/dryad.k67j5).

Goodman, R., Phillips, O. & Baker, T. (2013) Data from: The importance of crown dimensions to improve tropical tree biomass estimates. *Ecological Applications*. doi:[doi:10.5061/dryad.p281g](https://doi.org/doi:10.5061/dryad.p281g).

Grime, J.P., Hodgson, J.G. & Hunt, R. (2014) *Comparative Plant Ecology: A Functional Approach to Common British Species*. Springer.

Grootemaat, S., Wright, I., van, P., Bodegom, Cornelissen, J. & Cornwell, W. (2015) Data from: Burn or rot: leaf traits explain why flammability and decomposability are decoupled across species. *Functional Ecology*. doi:[doi:10.5061/dryad.m41f1](https://doi.org/doi:10.5061/dryad.m41f1).

Hamann, E., Kesselring, H., Armbruster, G., Scheepens, J. & StA?cklin, J. (2016) Data from: Evidence of local adaptation to fine- and coarse-grained environmental variability in Poa alpina in the Swiss Alps. *Journal of Ecology*. doi:[doi:10.5061/dryad.pt7n3](https://doi.org/doi:10.5061/dryad.pt7n3).

Hamilton, J., Lexer, C. & Aitken, S. (2012) Data from: Genomic and phenotypic architecture of a spruce hybrid zone (Picea sitchensis x P. glauca). *Molecular Ecology*. doi:[doi:10.5061/dryad.s11b6](https://doi.org/doi:10.5061/dryad.s11b6).

He, J.-S., Wang, Z., Wang, X., Schmid, B., Zuo, W., Zhou, M., Zheng, C., Wang, M. & Fang, J. (2006) A test of the generality of leaf trait relationships on the Tibetan Plateau. *New Phytologist*. 170 (4), 835–848.

Ishizuka, W. & Goto, S. (2011) Data from: Modeling intraspecific adaptation of Abies sachalinensis to local altitude and responses to global warming, based on a 36- year reciprocal transplant experiment. *Evolutionary Applications*. doi:[doi:10.5061/dryad.hh2g4s48](https://doi.org/doi:10.5061/dryad.hh2g4s48).

Karagatzides, J. & Ellison, A. (2008) Construction Costs of Carnivorous Plants and Non-Carnivorous Plants. *Harvard Forest Data Archive: HF112*.

King, D.A. (1996) Allometry and life history of tropical trees. *Journal of tropical ecology*. 12 (01), 25–44.

Kleinschroth, F., Healey, J., Sist, P., Mortier, F. & Go urlet-Fleury, S. (2016) Data from: How persistent are the impacts of logging roads on Central African forest vegetation? *Journal of Applied Ecology*. doi:[doi:10.5061/dryad.51p4f](https://doi.org/doi:10.5061/dryad.51p4f).

Kleyer, M., Bekker, R., Knevel, I., Bakker, J., Thompson, K., Sonnenschein, M., Poschlod, P., Van Groenendael, J., Klimeš, L., Klimešová, J., & others (2008) The LEDA Traitbase: a database of life-history traits of the Northwest European flora. *Journal of Ecology*. 96 (6), 1266–1274.

Kraft, N.J., Valencia, R. & Ackerly, D.D. (2008) Functional traits and niche-based tree community assembly in an Amazonian forest. *Science*. 322 (5901), 580–582.

Kraft, T., Wright, S., Turner, I., Lucas, P., Oufiero, C., Noor, M., Sun, I. & Dominy, N. (2015) Data from: Seed size and the evolution of leaf defences. *Journal of Ecology*. doi:[doi:10.5061/dryad.69ph0](https://doi.org/doi:10.5061/dryad.69ph0).

de La Riva, E., Pérez-Ramos, I., Tosto, A., Navarro-Fernández, C., Olmo, M., Marañón, T. & Villar, R. (2015) Data from: Disentangling the relative importance of species occurrence, abundance and intraspecific variability in community assembly: a trait-based approach at the whole-plant level in Mediterranean forests. *Oikos*. doi:[doi:10.5061/dryad.dr275.2](https://doi.org/doi:10.5061/dryad.dr275.2).

Lankinen, A. & Hydbom, S. (2016) Data from: Effects of soil resources on expression of a sexual conflict over timing of stigma receptivity in a mixed-mating plant. *Oikos*. doi:[doi:10.5061/dryad.2598k](https://doi.org/doi:10.5061/dryad.2598k).

Leishman, M., Cooke, J. & Richardson, D. (2014) Data from: Evidence for shifts to faster growth strategies in the new ranges of invasive alien plants. *Journal of Ecology*. doi:[doi:10.5061/dryad.2dj32](https://doi.org/doi:10.5061/dryad.2dj32).

Letcher, S., Lasky, J., Chazdon, R., Norden, N., Wright, S., et al. (2015) Data from: Environmental gradients and the evolution of successional habitat specialization: a test case with 14 Neotropical forest sites. *Journal of Ecology*. doi:[doi:10.5061/dryad.d87v7](https://doi.org/doi:10.5061/dryad.d87v7).

Li, W., Xu, F., Zheng, S., Taube, F. & Bai, Y. (2016a) Data from: Patterns and thresholds of grazing-induced changes in community structure and ecosystem functioning: species-level responses and the critical role of species traits. *Journal of Applied Ecology*. doi:[doi:10.5061/dryad.9n859](https://doi.org/doi:10.5061/dryad.9n859).

Li, X., Schmid, B., Wang, F. & Paine, C. (2016b) Data from: Net assimilation rate determines the growth rates of 14 species of subtropical forest trees. *PLOS ONE*. doi:[doi:10.5061/dryad.5kb61](https://doi.org/doi:10.5061/dryad.5kb61).

Liu, K., Eastwood, R., Flynn, S., Turner, R. & Stuppy, W. (2008) Seed information database (release 7.1, May 2008). *Available at ht tp://www. kew. org/data/sid*.

Louda, S.M., Dixon, P.M. & Huntly, N.J. (1987) Herbivory in sun versus shade at a natural meadow-woodland ecotone in the Rocky Mountains. *Plant Ecology*. 72 (3), 141–149.

Loughnan, D. & Gilbert, B. (2017) Data from: Trait-mediated community assembly: distinguishing the signatures of biotic and abiotic filters. *Oikos*. doi:[doi:10.5061/dryad.512p5](https://doi.org/doi:10.5061/dryad.512p5).

Maire, V., Wright, I., Prentice, I., Batjes, N., Bhaskar, R., van, P., Bodegom, Cornwell, W., Ellsworth, D., Niinemets, A., OrdoA?ez, A., Reich, P. & Santiago, L. (2015) Data from: Global effects of soil and climate on leaf photosynthetic traits and rates. *Global Ecology and Biogeography*. doi:[doi:10.5061/dryad.j42m7.2](https://doi.org/doi:10.5061/dryad.j42m7.2).

Manzano-Piedras, E., Marcer, A., Alonso-Blanco, C. & Picó, F. (2014) Data from: Deciphering the adjustment between environment and life history in annuals: lessons from a geographically-explicit approach in Arabidopsis thaliana. *PLoS ONE*. doi:[doi:10.5061/dryad.6nv8d](https://doi.org/doi:10.5061/dryad.6nv8d).

Martin, A., Rapidel, B., Roupsard, O., Van, K., den, Meersche, de, E., Melo, Virginio, Filho, Barrios, M. & Isaac, M. (2016) Data from: Intraspecific trait variation across multiple scales: the leaf economics spectrum in coffee. *Functional Ecology*. doi:[doi:10.5061/dryad.4t3r6](https://doi.org/doi:10.5061/dryad.4t3r6).

Marx, H., Giblin, D., Dunwiddie, P. & Tank, D. (2015) Data from: Deconstructing DarwinA?s naturalization conundrum in the San Juan Islands using community phylogenetics and functional traits. *Diversity and Distributions*. doi:[doi:10.5061/dryad.m88g7](https://doi.org/doi:10.5061/dryad.m88g7).

Mason, C. & Donovan, L. (2015) Data from: Evolution of the leaf economics spectrum in herbs: evidence from environmental divergences in leaf physiology across Helianthus (Asteraceae). *Evolution*. doi:[doi:10.5061/dryad.110s9](https://doi.org/doi:10.5061/dryad.110s9).

Mason, C., Goolsby, E., Humphreys, D. & Donovan, L. (2015) Data from: Phylogenetic structural equation modelling reveals no need for an ‘origin’ of the leaf economics spectrum. *Ecology Letters*. doi:[doi:10.5061/dryad.s652h](https://doi.org/doi:10.5061/dryad.s652h).

Mazer, S.J. (1989) Ecological, taxonomic, and life history correlates of seed mass among Indiana dune angiosperms. *Ecological Monographs*. 59 (2), 153–175.

McHugh, N., Edmondson, J., Gaston, K., Leake, J. & O’Sullivan, O. (2015) Data from: Modelling short-rotation coppice and tree planting for urban carbon management: a city-wide analysis. *Journal of Applied Ecology*. doi:[doi:10.5061/dryad.j25t0](https://doi.org/doi:10.5061/dryad.j25t0).

Medrano, M., Herrera, C. & Bazaga, P. (2014) Data from: Epigenetic variation predicts regional and local intraspecific functional diversity in a perennial herb. *Molecular Ecology*. doi:[doi:10.5061/dryad.fr2k8](https://doi.org/doi:10.5061/dryad.fr2k8).

Meers, T.L., Kasel, S., Bell, T.L. & Enright, N.J. (2010) Conversion of native forest to exotic Pinus radiata plantation: Response of understorey plant composition using a plant functional trait approach. *Forest Ecology and Management*. 259 (3), 399–409.

Milla, R., Morente-López, J., Alonso-Rodrigo, J., Martín-Robles, N. & Chapin, F., III (2014) Data from: Shifts and disruptions in resource-use trait syndromes during the evolution of herbaceous crops. *Proceedings of the Royal Society B*. doi:[doi:10.5061/dryad.dg85v](https://doi.org/doi:10.5061/dryad.dg85v).

Molinari, N. & D’Antonio, C. (2013) Data from: Structural, compositional and trait differences between native and non-native dominated grassland patches. *Functional Ecology*. doi:[doi:10.5061/dryad.50hd2](https://doi.org/doi:10.5061/dryad.50hd2).

Mottet, M., DeBlois, J. & Perron, M. (2015) Data from: High genetic variation and moderate to high values for genetic parameters of Picea abies resistance to Pissodes strobi. *Tree Genetics & Genomes*. doi:[doi:10.5061/dryad.pq075](https://doi.org/doi:10.5061/dryad.pq075).

Muir, C., Conesa, M., Roldán, E., Molins, A. & Galmés, J. (2016) Data from: Weak coordination between leaf structure and function among closely related tomato species. *New Phytologist*. doi:[doi:10.5061/dryad.1r8c2](https://doi.org/doi:10.5061/dryad.1r8c2).

Murali, K. (1997) Patterns of Seed Size, Germination and Seed Viability of Tropical Tree Species in Southern India. *Biotropica*. 29 (3), 271–279.

Neba, G., Newbery, D. & Chuyong, G. (2015) Data from: Limitation of seedling growth by potassium and magnesium supply for two ectomycorrhizal tree species of a Central African rain forest and its implication for their recruitment. *Ecology and Evolution*. doi:[doi:10.5061/dryad.k7729](https://doi.org/doi:10.5061/dryad.k7729).

Nidzgorski, D. & Hobbie, S. (2016) Data from: Urban trees reduce nutrient leaching to groundwater. *Ecological Applications*. doi:[doi:10.5061/dryad.n3s2m](https://doi.org/doi:10.5061/dryad.n3s2m).

Niu, K., He, J. & Lechowicz, M. (2016) Data from: Grazing-induced shifts in community functional composition and soil nutrient availability in Tibetan alpine meadows. *Journal of Applied Ecology*. doi:[doi:10.5061/dryad.r5m20](https://doi.org/doi:10.5061/dryad.r5m20).

Norghauer, J., Glauser, G. & Newbery, D. (2014) Data from: Seedling resistance, tolerance and escape from herbivores: insights from co-dominant canopy tree species in a resource-poor African rain forest. *Functional Ecology*. doi:[doi:10.5061/dryad.129vg](https://doi.org/doi:10.5061/dryad.129vg).

Ostevik, K., Andrew, R., Otto, S. & Rieseberg, L. (2016) Data from: Multiple reproductive barriers separate recently diverged sunflower ecotypes. *Evolution*. doi:[doi:10.5061/dryad.223p4](https://doi.org/doi:10.5061/dryad.223p4).

Osuri, A. & Sankaran, M. (2016) Data from: Seed size predicts community composition and carbon storage potential of tree communities in rainforest fragments in IndiaA?s Western Ghats. *Journal of Applied Ecology*. doi:[doi:10.5061/dryad.7s7r1](https://doi.org/doi:10.5061/dryad.7s7r1).

Paine, C., Amissah, L., Auge, H., Baraloto, C., Baruffol, M., et al. (2015) Data from: Globally, functional traits are weak predictors of juvenile tree growth, and we do not know why. *Journal of Ecology*. doi:[doi:10.5061/dryad.h9083](https://doi.org/doi:10.5061/dryad.h9083).

Paynter, Q., Buckley, Y., Peterson, P., Go urlay, A. & Fowler, S. (2015) Data from: Breaking and remaking a seed and seed predator interaction in the introduced range of Scotch Broom (Cytisus scoparius) in New Zealand. *Journal of Ecology*. doi:[doi:10.5061/dryad.dd3ph.2](https://doi.org/doi:10.5061/dryad.dd3ph.2).

Peet, R.K., Lee, M.T., Boyle, M.F., Wentworth, T.R., Schafale, M.P. & Weakley, A.S. (2012) Vegetation-plot database of the Carolina Vegetation Survey. *Biodiversity and Ecology*. 4, 243–253.

Pérez-de-Lis, G., Olano, J., Rozas, V., Rossi, S., Vázquez-Ruiz, R. & García-González, I. (2016) Data from: Environmental conditions and vascular cambium regulate carbon allocation to xylem growth in deciduous oaks. *Functional Ecology*. doi:[doi:10.5061/dryad.1cn19](https://doi.org/doi:10.5061/dryad.1cn19).

van der Plas, F., Howison, R., Mpanza, N., Cromsigt, J. & Olff, H. (2016) Data from: Different-sized grazers have distinctive effects on plant functional composition of an African savannah. *Journal of Ecology*. doi:[doi:10.5061/dryad.512m0](https://doi.org/doi:10.5061/dryad.512m0).

Ploton, P., Barbier, N., Momo, S., Réjou-Méchain, M., Boyemba, F., Bosela, et al. (2016) Data from: Closing a gap in tropical forest biomass estimation: taking crown mass variation into account in pantropical allometries. *Biogeosciences*. doi:[doi:10.5061/dryad.f2b52](https://doi.org/doi:10.5061/dryad.f2b52).

Poorter, L. (2008) The Relationships of Wood-, Gas- and Water Fractions of Tree Stems to Performance and Life History Variation in Tropical Trees. *Annals of Botany*. 102 (3), 367. doi:[10.1093/aob/mcn103](https://doi.org/10.1093/aob/mcn103).

Poorter, L. & Bongers, F. (2006) Leaf traits are good predictors of plant performance across 53 rain forest species. *Ecology*. 87 (7), 1733–1743.

Poorter, L. & Rozendaal, D.M. (2008) Leaf size and leaf display of thirty-eight tropical tree species. *Oecologia*. 158 (1), 35–46.

Price, C., Wright, I., Ackerly, D., Niinemets, Ü., Reich, P. & Veneklaas, E. (2014) Data from: Are leaf functional traits ‘invariant’ with plant size, and what is ‘invariance’ anyway? *Functional Ecology*. doi:[doi:10.5061/dryad.r3n45](https://doi.org/doi:10.5061/dryad.r3n45).

Ramirez-Valiente, J., Lorenzo, Z., Soto, A., Valladares, F., Gil, L. & Aranda, I. (2009) Data from: Elucidating the role of genetic drift and natural selection in cork oak differentiation regarding drought tolerance. *Molecular Ecology*. doi:[doi:10.5061/dryad.1284](https://doi.org/doi:10.5061/dryad.1284).

Rasmann, S. & Agrawal, A. (2011) Data from: Evolution of specialization: a phylogenetic study of host range in the red milkweed beetle (Tetraopes tetraophthalmus). *The American Naturalist*. doi:[doi:10.5061/dryad.8557](https://doi.org/doi:10.5061/dryad.8557).

Robinson, K., Hauzy, C., Loeuille, N. & Albrectsen, B. (2015) Data from: Relative impacts of environmental variation and evolutionary history on the nestedness and modularity of tree-herbivore networks. *Ecology and Evolution*. doi:[doi:10.5061/dryad.4q78p](https://doi.org/doi:10.5061/dryad.4q78p).

Rodríguez-Quilón, I., Santos-del-Blanco, L., Serra-Varela, M., Koskela, J., González-Martínez, S. & Alía, R. (2016) Data from: Capturing neutral and adaptive genetic diversity for conservation in a highly structured tree species. *Ecological Applications*. doi:[doi:10.5061/dryad.c289v](https://doi.org/doi:10.5061/dryad.c289v).

Roe, A., MacQuarrie, C., Gros-Louis, M., Simpson, J., Lamarche, J., Beardmore, T., Thompson, S., Tanguay, P. & Isabel, N. (2014) Data from: Fitness dynamics within a poplar hybrid zone: II. Impact of exotic sex on native poplars in an urban jungle. *Ecology and Evolution*. doi:[doi:10.5061/dryad.6vk6f](https://doi.org/doi:10.5061/dryad.6vk6f).

Royer, D.L., Wilf, P., Janesko, D.A., Kowalski, E.A. & Dilcher, D.L. (2005) Correlations of climate and plant ecology to leaf size and shape: potential proxies for the fossil record. *American Journal of Botany*. 92 (7), 1141–1151.

Russo, S.E., Jenkins, K.L., Wiser, S.K., Uriarte, M., Duncan, R.P. & Coomes, D.A. (2010) Interspecific relationships among growth, mortality and xylem traits of woody species from New Zealand. *Functional Ecology*. 24 (2), 253–262.

Salgado-Luarte, C. & Gianoli, E. (2012) Data from: Herbivores modify selection on plant functional traits in a temperate rainforest understory. *The American Naturalist*. doi:[doi:10.5061/dryad.53tr05j2](https://doi.org/doi:10.5061/dryad.53tr05j2).

Sánchez-Robles, J., García Castaño, J., Balao, F., Terrab, A., Navarro, L., Sampedro, Tremetsberger, K. & Talavera, S. (2014) Data from: Effects of tree architecture on pollen dispersal and mating patterns in Abies pinsapo Boiss. (Pinaceae). *Molecular Ecology*. doi:[doi:10.5061/dryad.f0d93](https://doi.org/doi:10.5061/dryad.f0d93).

Schneider, G., Krauss, J., Riedinger, V., Holzschuh, A. & Steffan-Dewenter, I. (2015) Data from: Biological pest control and yields depend on spatial and temporal crop cover dynamics. *Journal of Applied Ecology*. doi:[doi:10.5061/dryad.q9690](https://doi.org/doi:10.5061/dryad.q9690).

Sessa, E. & Givnish, T. (2013) Data from: Leaf form and photosynthetic physiology of Dryopteris species distributed along light gradients in eastern North America. *Functional Ecology*. doi:[doi:10.5061/dryad.38h06](https://doi.org/doi:10.5061/dryad.38h06).

Shibata, R., Kurokawa, H., Shibata, M., Tanaka, H., Iida, S., Masaki, T. & Nakashizuka, T. (2015) Data from: Relationships between resprouting ability, species traits, and resource allocation patterns in woody species in a temperate forest. *Functional Ecology*. doi:[doi:10.5061/dryad.rj480](https://doi.org/doi:10.5061/dryad.rj480).

Shugart Jr, H., Hopkins, M., Burgess, I. & Mortlock, A. (1980) Development of a succession model for subtropical rain forest and its application to assess the effects of timber harvest at Wiangaree State Forest, New South Wales. *J. Environ. Manage.;(United States)*. 11 (3).

Simpson, K., Ripley, B., Christin, P., Belcher, C., Lehmann, C., Thomas, G. & Osborne, C. (2015) Data from: Determinants of flammability in savanna grass species. *Journal of Ecology*. doi:[doi:10.5061/dryad.2c506](https://doi.org/doi:10.5061/dryad.2c506).

Spasojevic, M., Turner, B. & Myers, J. (2015) Data from: When does intraspecific trait variation contribute to functional beta-diversity? *Journal of Ecology*. doi:[doi:10.5061/dryad.rr4pm](https://doi.org/doi:10.5061/dryad.rr4pm).

Steane, D., Potts, B., McLean, E., Collins, L., Prober, S., Stock, W., Vaillancourt, R. & Byrne, M. (2015) Data from: Genome-wide scans reveal cryptic population structure in a dry-adapted eucalypt. *Tree Genetics & Genomes*. doi:[doi:10.5061/dryad.h06r3](https://doi.org/doi:10.5061/dryad.h06r3).

Steane, D., Potts, B., McLean, E., Prober, S., Stock, W., Vaillancourt, R. & Byrne, M. (2014) Data from: Genome-wide scans detect adaptation to aridity in a widespread forest tree species. *Molecular Ecology*. doi:[doi:10.5061/dryad.qq20s](https://doi.org/doi:10.5061/dryad.qq20s).

Stevens, J., Safford, H., Harrison, S. & Latimer, A. (2015) Data from: Forest disturbance accelerates thermophilization of understory plant communities. *Journal of Ecology*. doi:[doi:10.5061/dryad.q2n8p](https://doi.org/doi:10.5061/dryad.q2n8p).

Storkey, J., DA?ring, T., Baddeley, J., Collins, R., Roderick, S., Jones, H. & Watson, C. (2014) Data from: Engineering a plant community to deliver multiple ecosystem services. *Ecological Applications*. doi:[doi:10.5061/dryad.qj3mg](https://doi.org/doi:10.5061/dryad.qj3mg).

Szefer, P., Carmona, C., Chmel, K., KonecnA!, M., Libra, M., Molem, K., Novotny, V., Segar, S., A?vamberkovA!, E., Topliceanu, T. & Leps, J. (2016) Data from: Determinants of litter decomposition rates in a tropical forest: functional traits, phylogeny and ecological succession. *Oikos*. doi:[doi:10.5061/dryad.4b95c.2](https://doi.org/doi:10.5061/dryad.4b95c.2).

Thomas, S., Martin, A. & Mycroft, E. (2015) Data from: Tropical trees in a wind-exposed island ecosystem: height-diameter allometry and size at onset of maturity. *Journal of Ecology*. doi:[doi:10.5061/dryad.bs332](https://doi.org/doi:10.5061/dryad.bs332).

Urrutia-Jalabert, R., Malhi, Y. & Lara, A. (2015) Data from: The oldest, slowest forests in the world? Exceptional biomass and slow carbon dynamics of Fitzroya cupressoides temperate rainforests in southern Chile. *PLOS ONE*. doi:[doi:10.5061/dryad.2kh91](https://doi.org/doi:10.5061/dryad.2kh91).

Vincent, J., Weiblen, G. & May, G. (2015) Data from: Host associations and beta diversity of fungal endophyte communities in New Guinea rainforest trees. *Molecular Ecology*. doi:[doi:10.5061/dryad.7q1b2](https://doi.org/doi:10.5061/dryad.7q1b2).

Welsh, M., Cronin, J. & Mitchell, C. (2016) Data from: The role of habitat filtering in the leaf economics spectrum and plant susceptibility to pathogen infection. *Journal of Ecology*. doi:[doi:10.5061/dryad.356v3](https://doi.org/doi:10.5061/dryad.356v3).

Wigley, B., Slingsby, J., Diaz, S., Bond, W., Fritz, H. & Coetsee, C. (2016) Data from: Leaf traits of African woody savanna species across climate and soil fertility gradients: evidence for conservative vs. acquisitive resource use strategies. *Journal of Ecology*. doi:[doi:10.5061/dryad.v240b](https://doi.org/doi:10.5061/dryad.v240b).

Wiser, S.K., Bellingham, P.J. & Burrows, L.E. (2001) Managing biodiversity information: development of New Zealand’s National Vegetation Survey databank. *N. Z. J. Ecol.* 25 (2), 1–17.

Wood, Z., Peart, D., Palmiotto, P., Kong, L. & Peart, N. (2015) Data from: Asymptotic allometry and transition to the canopy in Abies balsamea. *Journal of Ecology*. doi:[doi:10.5061/dryad.r3645](https://doi.org/doi:10.5061/dryad.r3645).

Wright, I.J., Reich, P.B., Westoby, M., Ackerly, D.D., Baruch, Z., Bongers, F., Cavender-Bares, J., Chapin, T., Cornelissen, J.H., Diemer, M., & others (2004) The worldwide leaf economics spectrum. *Nature*. 428 (6985), 821–827.

Yang, X., Xia, H., Wang, W., Wang, F., Su, J., Snow, A. & Lu, B. (2011) Data from: Transgenes for insect resistance reduce herbivory and enhance fecundity in advanced generations of crop-weed hybrids of rice. *Evolutionary Applications*. doi:[doi:10.5061/dryad.8974](https://doi.org/doi:10.5061/dryad.8974).

Yoder, J., Smith, C., Rowley, D., Flatz, R., Godsoe, W., Drummond, C. & Pellmyr, O. (2013) Data from: Effects of gene flow on phenotype matching between two varieties of Joshua tree (Yucca brevifolia; Agavaceae) and their pollinators. *Journal of Evolutionary Biology*. doi:[doi:10.5061/dryad.369q9](https://doi.org/doi:10.5061/dryad.369q9).

Zanne, A., Oberle, B., Dunham, K., Milo, A., Walton, M. & Young, D. (2015) Data from: A deteriorating state of affairs: how endogenous and exogenous factors determine plant decay rates. *Journal of Ecology*. doi:[doi:10.5061/dryad.b3q08](https://doi.org/doi:10.5061/dryad.b3q08).

Zas, R., CendA!n, C. & Sampedro, L. (2013) Data from: Mediation of seed provisioning in the transmission of environmental maternal effects in Maritime pine (Pinus pinaster Aiton). *Heredity*. doi:[doi:10.5061/dryad.k7c1m](https://doi.org/doi:10.5061/dryad.k7c1m).

Zas, R. & Sampedro, L. (2014) Data from: Heritability of seed weight in Maritime pine, a relevant trait in the transmission of environmental maternal effects. *Heredity*. doi:[doi:10.5061/dryad.60j76](https://doi.org/doi:10.5061/dryad.60j76).

Zerebecki, R., Crutsinger, G. & Hughes, A. (2016) Data from: Spartina alterniflora genotypic identity affects plant and consumer responses in an experimental marsh community. *Journal of Ecology*. doi:[doi:10.5061/dryad.h32p6](https://doi.org/doi:10.5061/dryad.h32p6).

Zheng, S., Ren, H., Lan, Z., Li, W., Wang, K. & Bai, Y. (2010) Effects of grazing on leaf traits and ecosystem functioning in Inner Mongolia grasslands: scaling from species to community. *Biogeosciences*. 7 (3), 1117–1132.

Zheng, Z., Zhang, S., Baskin, C., Baskin, J., Schaefer, D., Yang, X. & Yang, L. (2015) Data from: Hollows in living trees develop slowly but considerably influence the estimate of forest biomass. *Functional Ecology*. doi:[doi:10.5061/dryad.dh11k](https://doi.org/doi:10.5061/dryad.dh11k).

Zhu, H., Fu, B., Wang, S., Zhu, L., Jiao, L. & Wang, C. (2015) Data from: Reducing soil erosion by improving community functional diversity in semi-arid grasslands. *Journal of Applied Ecology*. doi:[doi:10.5061/dryad.b5tr9](https://doi.org/doi:10.5061/dryad.b5tr9).

Zuppinger-Dingley, D., Schmid, B., Petermann, J., Yadav, V., De, G., Deyn & Flynn, D. (2014) Data from: Selection for niche differentiation in mixed plant communities increases biodiversity effects. *Nature*. doi:[doi:10.5061/dryad.750df](https://doi.org/doi:10.5061/dryad.750df).

Züst, T. & Agrawal, A. (2015) Data from: Population growth and sequestration of plant toxins along a gradient of specialization in four aphid species on the common milkweed Asclepias syriaca. *Functional Ecology*. doi:[doi:10.5061/dryad.35nn0](https://doi.org/doi:10.5061/dryad.35nn0).
